# Transposon end recognition and pairing by I-F3 CRISPR-associated transposase

**DOI:** 10.64898/2026.07.09.737429

**Authors:** Vinh H. Truong, Darcie J. Miller, Shirin Fatma, Yuewen Sheng, Chinmai Pindi, Mohd Ahsan, Giulia Palermo, Elizabeth H. Kellogg

## Abstract

To develop gene therapy tools based on CRISPR-associated transposons (CASTs), it is essential to define how transposon ends are recognized and paired during transposition. Tn7-like transposons typically contain asymmetric left- and right-end sequences that flank and define DNA cargo. However, how the transposase recognizes these different sequences and assembles them into a paired end complex for cut-and-paste transposition remains unknown. Here we present the cryo-EM structure of type I-F3 (VchCAST) CAST transposase TnsB in complex with transposon DNA ends and host factor IHF, along with biochemistry, molecular dynamics, and in vivo analyses. Our structure reveals the stoichiometry and architecture of the assembly, as well as the DNA distortions required to accommodate transposon end asymmetry. Molecular dynamics suggests that these distortions are required for the coordinated assembly of the complex. Physical pairing of asymmetric left and right ends results in a novel protein-protein interface that is required for transposition efficiency. Our findings explain how transposases regulate the pairing of transposon end sequences for high-fidelity recognition, reveal a model of transposon end synapsis, and suggest future avenues to engineer DNA cargo for genome-editing applications.

## Introduction

Transposons, or ‘jumping genes’, are segments of DNA that autonomously mobilize within a host genome. Class II DDE transposons encode a transposase that catalyzes excision and genomic integration of transposon DNA in a process known as cut-and-paste transposition^1^. Transposition proceeds as follows: the transposase brings together, or synapses, transposon right and left end sequences and, through a multi-step process, catalyzes the strand-esterification reaction to integrate donor DNA into the genome (see Arinkin et al. for a detailed review)^1^. Importantly, transposon genomic integration does not require introduction of cytotoxic double-stranded DNA breaks.

Transposon end sequences can be transferred onto exogenous donor DNA to direct integration of large DNA cargo into the genome, making transposons attractive and highly adaptable as genome-editing tools. These end sequences can be hundreds of base-pairs long, placing considerable restrictions on donor DNA cargo sequences. Transposons such as sleeping beauty^2^, IstA/B^3^, and Mos1^4^ have symmetric ends. In contrast, transposons with asymmetric ends can have different number, orientation, and spacing of transposase binding sites in the right and left ends^5,6^. This asymmetry provides additional regulatory control over transposition, enabling the transposase to maintain orientation of donor DNA insertions. This is presumed to occur through a higher order synaptic complex, but there are few structures showing how asymmetric ends might be accommodated in such an assembly^5,6^.

Much more structural information is available for the catalytic core of DDE transposases^7^, similar to retroviral integrases^8^, which assemble into a strikingly consistent, characteristic two-fold symmetric complex poised to integrate donor DNA^9^. End binding and catalysis occurs in *trans*: each half of the assembly engages with one end of the donor DNA while positioning the RNase H catalytic domain to act on the other end^9^. In the case of Tn7-like transposons^10^ and its distant homologs such as MuA^11^, the catalytic core corresponds to a two-fold symmetric homo-tetramer that, despite evolutionary distance, retains an essentially identical architecture. In some Tn7-like transposons, an additional endonuclease, TnsA, associates with the transposase^12^. TnsA cuts the 5’ end of transposon DNA to create simple insertions rather than co-integrates^13^ and potentially stimulates TnsB DNA-binding^12^.

Tn7-like transposons have the remarkable property of being site specific^14^ due to the coordinated function of their transposase (TnsB) with other transposon-encoded proteins^10^. The Tn7-like superfamily has diverse transposon end configurations^15^. Tn7 itself has four partially overlapping TnsB-binding sites in its right end and three well-separated sites in its left end^16^. A recent cryo-EM structure of TnsB bound to its right end DNA reveals a tiled oligomeric assembly that differs from the characteristic catalytic core structure^17^, suggesting that different assemblies may form during earlier steps of transposition. However, the mechanism by which TnsB recognizes and pairs distinct transposon end sequences with such high fidelity during the initial stages of transposition remains an open question.

CRISPR-associated transposons, or CASTs, are recently discovered Tn7-like transposons that rely on a CRISPR-like effector and a homolog of TnsD, called TniQ, rather than a sequence-specific DNA-binding protein^18^. CASTs are an exciting class of Tn7-like transposons because they can be developed into versatile genome-editing tools, capable of integrating large DNA cargo up to 10 Kb in size with precise spacing and orientation in a guide-RNA dependent manner^19,20^. The I-F3 CAST family has notable on-target integration specificity^21^, which has proven useful for applications such as microbial strain engineering^22^ and, potentially, clinical gene therapy^23,24^. It is essential to understand how the transposon end sequences are recognized and paired by the CAST transposase: such knowledge would allow us to modify transposon end sequences without affecting transposition activity, thus removing limitations for cargo DNA sequences.

Here we set out to determine how TnsB from I-F3 CAST (VchCAST or Tn6677) recognizes and pairs its asymmetric transposon ends^14^, which is required for on-target donor DNA integration^25^. This transposon has three 20 base-pair long TnsB-binding sites unevenly spaced in its left end (LE1–3) and three on its right end (RE1–3) (**Figure 1A**); all except RE3 are required for activity^19^. In addition, there is a conserved IHF-binding site between LE1 and LE2^25^; IHF is a host factor that stimulates transposition by an order of magnitude (**Figure 1A**), by unknown mechanisms. We present structural, biochemistry, molecular dynamics simulations, and *in vivo* analyses that provide insights into how the initial stages of synaptic complex formation occur, revealing dramatic DNA distortions. Further, we identify a stabilizing protein-protein interface that only occurs when both left and right transposon ends are present within the assembly, providing a compelling mechanistic explanation for how transposon end synapsis can occur with high fidelity.

**Figure 1.**
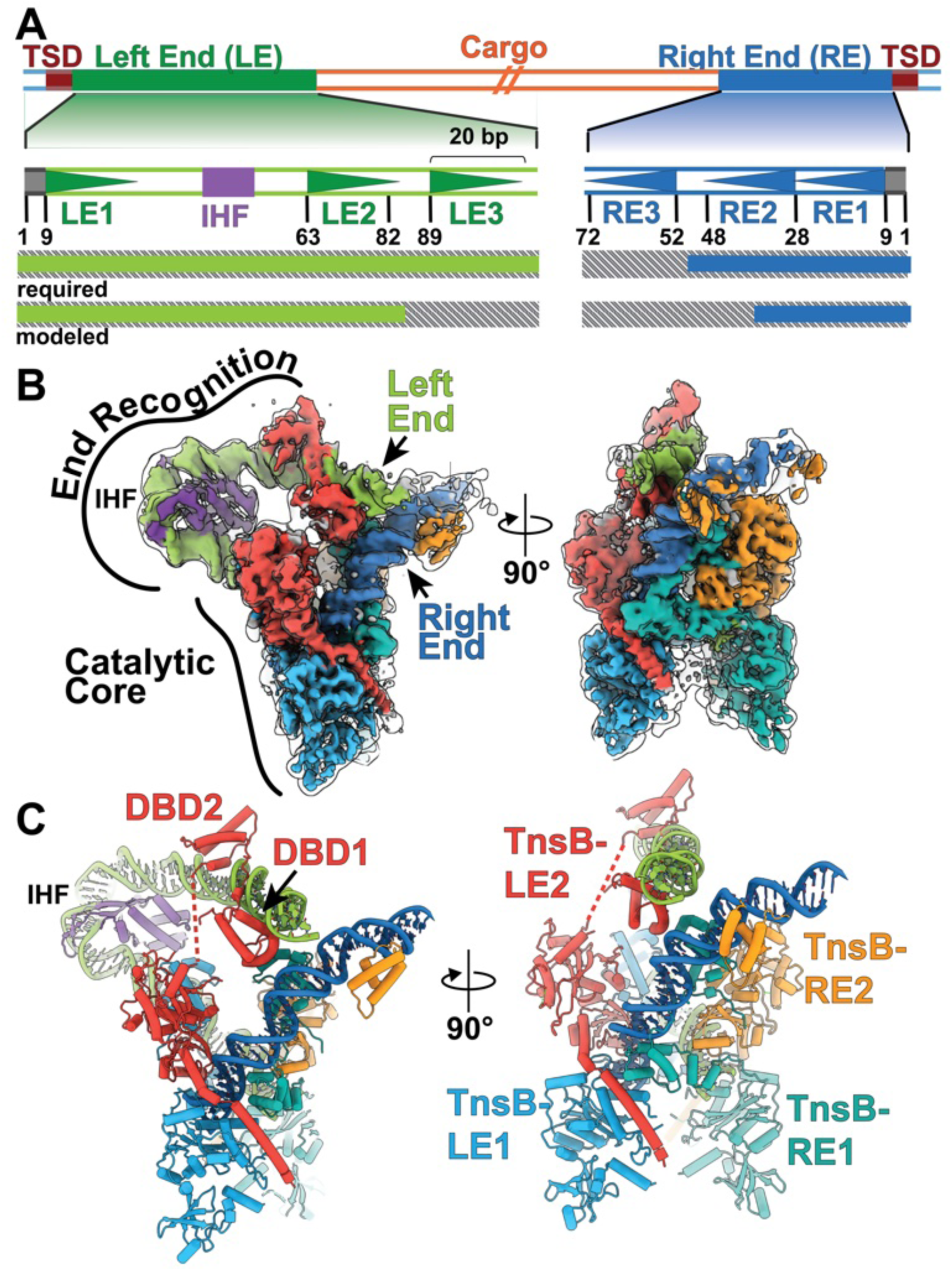
Architecture of the VchCAST transposase with paired transposon ends. **A.** VchCAST transposon end organization, with left end in green and right end in blue. The target-site duplication (TSD, in brown) is not considered part of the transposon. Cargo (orange) can be of variable size. Transposase binding sites are numbered starting from the site closest to the TSD (LE1–LE3 and RE1–RE3); triangles show orientation, and numbers below indicate base-pair positions. Gray boxes at each far end indicates the 8-basepair terminal sequence required for catalysis. Purple box, IHF binding site. Bottom, bars show portions of the transposon ends that are required for integration and that are modeled in our cryo-EM structure. **B.** Final composite map, shown in solid surface (displayed at threshold = 9.24), and locally filtered consensus map, shown as transparent outline (displayed at threshold = 0.08). **C.** Atomic model shown in cartoon representation. In panels b and c, transposon ends are colored as in panel A. IHF is colored purple. Transposase subunits are labeled in panel c according to the transposase binding site they recognize: TnsB-LE1 (light blue), TnsB-LE2 (red), TnsB-RE1 (aqua), TnsB-RE2 (yellow). DNA-binding domains, DBD1 (residues 37–113) and DBD2 (residues 125–184) are labeled for clarity in TnsB-LE2. Dotted lines indicate unmodeled regions.

## Results

### Asymmetric architecture of transposon end synaptic complex

We reconstituted a nucleoprotein complex with TnsB, IHF, and full left and right transposon ends, using our previously established strategy^26^. Briefly, we designed a strand-transfer substrate corresponding to a branched Shapiro intermediate, mimicking the integration product (**Figure S1**). We used single-particle cryo-EM to obtain a 3.9 Å-resolution structure of an assembly containing 4 TnsB subunits and 1 IHF dimer (**Figure 1B, Table 1**). The local resolution ranges between ∼3.7–10 Å resolution, due to the intrinsic flexibility of the long DNA substrates (**Figure S2, Movie S1**). We performed local refinement of specific regions (**Figure S3**) and used the locally refined maps to create a composite map, which was used for model refinement. We assigned the majority of each of the four TnsB subunits by docking in portions of AlphaFold models^27^ (**Figure 1C**), and named each according to the transposase binding site they bind (hence, TnsB-RE1 and so on) (**Figure 1C**).

The tetrameric structure has two distinct halves: an asymmetric half (here named end recognition) associates with RE2 and LE2 (**Figure 1B**), whereas the other half has two-fold symmetry and corresponds to the catalytic core (**Figure 1B**), adopting an architecture resembling other strand-transfer complex (STC) structures^26,11^. Like other STC structures, the catalytic core is homogeneously high resolution, with clear density for protein sidechains and for DNA phosphate backbone and bases (**Figure S4**). IHF causes a 180° bend in the DNA (**Figure S5**), positioning LE2 to interact with the N-terminal DNA-binding domains (DBD1 and DBD2) of TnsB-LE2 (**Figure 1C**). This dramatic distortion is required for LE2 to be accessible to the protein. Moreover, the spatial arrangement and phasing of LE2 within the structure suggest that its recognition by TnsB is highly sensitive to its position, which is consistent with the previously observed 10-base-pair periodicity in transposition efficiency when the distance between LEs was altered^25^. We observe DBD1 of TnsB-RE2 bound to RE2 in the cryo-EM map, but given the limited local resolution (8.0–9.0 Å) (**Figure S2**) and its small size (77 amino acids) (**Figure S6**), we did not model this region.

Since we supplied a strand-transfer DNA substrate, we expected to see target-DNA density, as in previous structures^26,5^, but that density was weak (**Figure S7**), and we were only able to reliably trace DNA up to the beginning of the target-site duplication. Thus, this structure’s features correspond more closely to a paired-end complex rather than a strand-transfer complex. It is possible that additional transposon-encoded factors, either TnsA or TnsC, known to interact with TnsB^12,28^, are required to stabilize target-DNA binding, and future work should identify additional regulatory mechanisms that contribute to the exquisite site-specificity of the I-F3 CAST subfamily.

In the donor DNA, we observe distortions in both right and left transposon ends that extend beyond those introduced by IHF binding. The left end DNA is bent approximately 34° between the IHF and LE2 (**Figure S8A-B**). This appears required for IHF and TnsB-LE2 to simultaneously occupy their respective sites, as modeling ideal B-form DNA results in steric clashes (**Figure S8A-B)**. Additionally, this distortion aligns well with the footprint of the bound DBD1 and DBD2. We compared our structure to a previous structure of the Tn7 transposase bound to a DNA substrate representing the full right end (PDB 7PIK)^17^. The N-terminal DNA-binding domains of Tn7 and I-F3 CAST transposases have similar folds and DNA-binding modes, but the path of the DNA molecule is substantially different (**Figure S8C**), suggesting that DNA distortions are dependent on the surrounding context, rather than the specifics of TnsB-DNA interactions.

### A new TnsB inter-subunit interface formed through IHF-induced DNA distortions

In our structure, the right and left ends are in close physical proximity, crossing over one another at the ends of LE2 and RE2 (**Figure 1C & S8A**). This arrangement appears to be mediated in part by interactions between TnsB subunits, specifically via the DBD2 domains from TnsB-LE2 and TnsB-RE1 (**Figure 2A**). This interaction is mediated by hydrophobic interactions involving V78, and mutations at this position substantially reduced integration activity (**Figure 2B**). In particular, mutations V78D and V78K caused a 10-fold reduction in transposition activity, similar to the effect of IHF deletion^25^. To understand the effect of these mutations on this assembly, we performed molecular dynamics (MD) simulations with the WT and mutant TnsB. Introduction of V78K mutation caused the TnsB-LE2 and TnsB-RE1 subunits to dissociate, driven by electrostatic repulsion between positively charged side chains on both domains (**Figure 2C-D**). This loss of interaction provides a direct mechanistic explanation for the reduced transposition activity. In contrast, the V78D mutation preserved local contacts between TnsB-LE2 and TnsB-RE1 through compensatory interactions with nearby lysine residues (**Figure S9**). To further delineate the structural consequences of these mutations, we examined broader interaction networks across the assembled complex. V78K selectively destabilized the TnsB-LE2 and TnsB-RE1 interface (**Figure 2E**), while V78D perturbed distal interfaces, particularly those between TnsB-LE2 and TnsB-LE1 (**Figure S10**). These observations suggest that V78K impairs function primarily by weakening the interface at the mutation site, while V78D exerts a more distributed destabilizing effect, implicating a distinct mechanism underlying its reduced activity.

**Figure 2.**
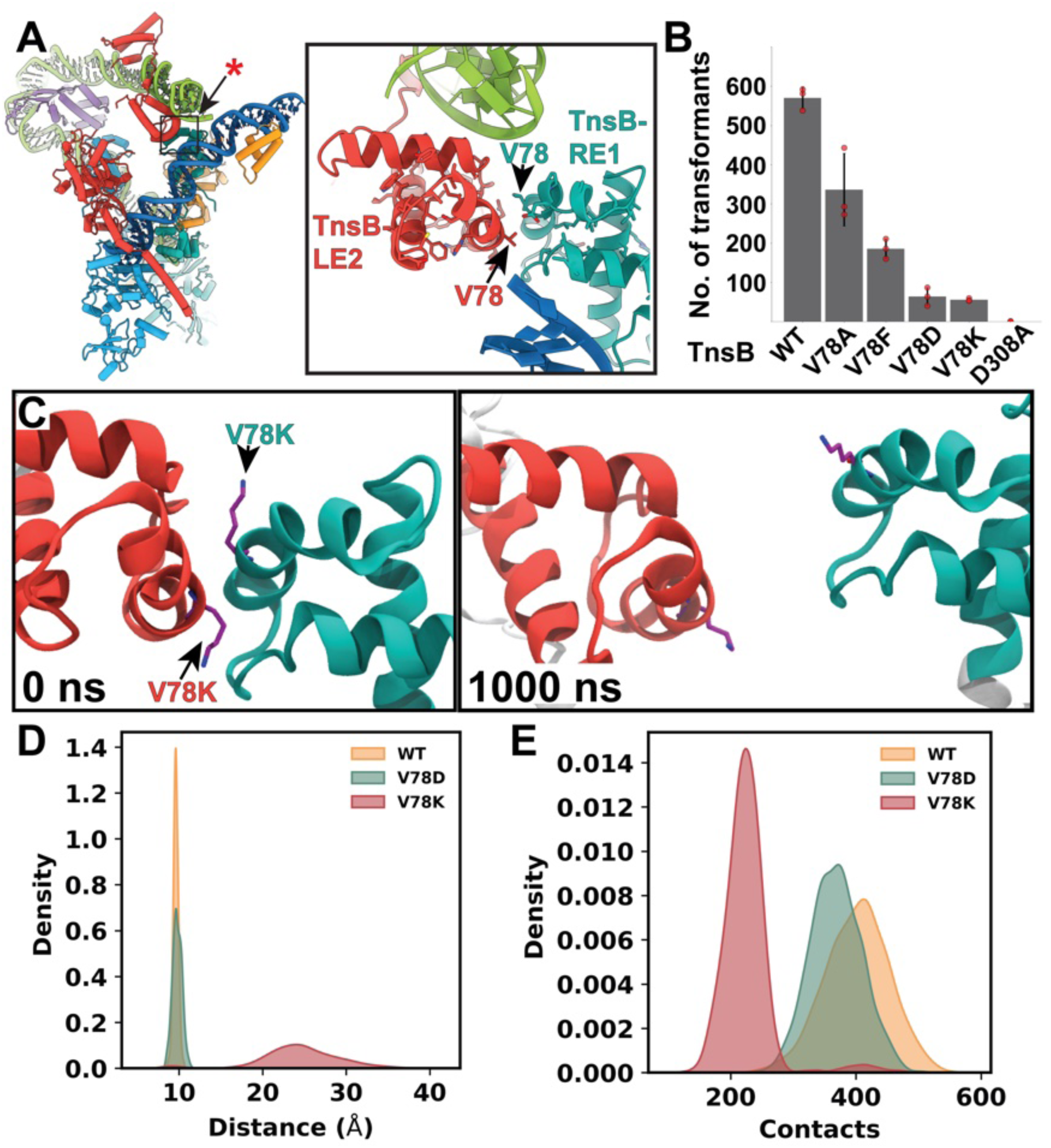
DNA distortions result in novel TnsB inter-subunit contacts that are important for transposition. **A.** Left, atomic model with same color scheme as in figure 1. Red asterisk indicates region of interest shown on the right, in the same viewing direction and highlighting TnsB inter-subunit contacts and showing Val78. **B.** In vivo transposition assay of TnsB V78 mutations. WT, wild-type. Red dots indicate biological triplicate measurements and black bars show standard error. **C.** Molecular dynamics (MD) simulation snapshots of the V78K mutant, showing electrostatic repulsion at the DBD2 interface, resulting in separation of TnsB-LE2 (red) and TnsB-RE1 (cyan). Timepoint in nanoseconds (ns) indicated at the bottom left. **D.** Distribution of distances between residues at position 78 on TnsB-LE2 and TnsB-RE1 from MD simulations for both V78K and V78D. **E** Density plots of inter-subunit contacts from MD simulations for both V78K and V78D, measured at the interface indicated with red asterisk in panel A.

We also reconstituted and imaged the complex in the absence of IHF: the 2D-class average reveals a 2-fold symmetric assembly (**Figure 3A**), consistent with a simulated symmetrized version of TnsB bound to two transposon right ends (**Figure 3B**). This observation is in line with data indicating that two right ends can still support reduced levels of transposition, but not two left ends^25^. To probe the functional role of IHF further, we performed MD simulations of the complex with and without IHF bound. In the absence of IHF, we observed increased flexibility in the DBD regions (residues 100-190) of the TnsB-LE1 subunit (**Figure 3C & Figure S11**), accompanied by reduced stability of the left-end DNA (**Figure 3D**). To understand the effect of IHF on DNA architecture, we also simulated the left-end DNA alone, with and without IHF. IHF maintains the sharply bent DNA conformation observed in the cryo-EM structure, whereas its removal leads to rapid DNA relaxation into a straight conformation (**Figure S12**). This demonstrates that IHF imposes and stabilizes the left-end DNA architecture necessary for productive assembly of the complex, and supports the experimental observation that IHF absence results in a different architecture (**Figure 3A-B**). Together, these structural, biochemical, and computational findings indicate that efficient formation of the observed TnsB–TnsB interface requires IHF-induced pre-organization of the left-end DNA into the bent conformation captured in our structure.

**Figure 3.**
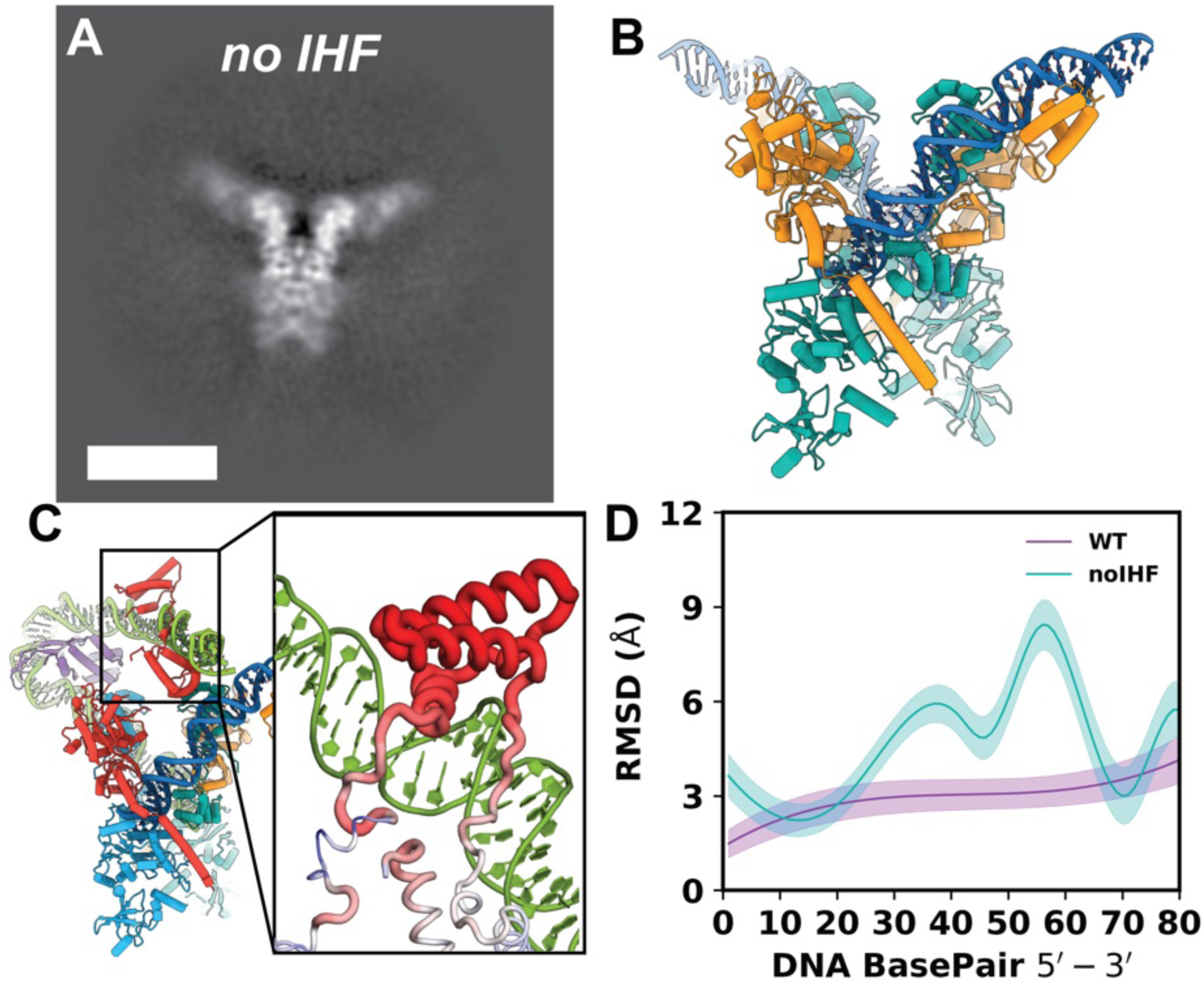
IHF is essential for inducing distortions and stabilizing the observed assembly. **A.** Representative cryo-EM 2D class average of VchCAST complex in the absence of IHF. Scale bar, 100 Å. **B.** Simulated complex of VchCAST containing 2 transposon right ends, generated by symmetrizing the right half of the VchCAST complex containing IHF. **C.** Flexibility of the VchCAST complex in the absence of IHF, annotated by per-residue root-mean-square fluctuations (RMSF) from molecular dynamics (MD) simulations. Regions of increased flexibility are shown as thicker tubes (red), highlighting pronounced mobility within the DBD2 domain of TnsB-LE1. **D.** Root-mean-square deviations (RMSD) of the left-end DNA backbone from MD simulations of the WT complex (purple) and the assembly lacking IHF (green). The absence of IHF destabilizes the left-end DNA, highlighting the base pairs with higher deviation.

### Base-specific interactions explain position-specific sequence constraints

Since transposases have remarkable specificity for their ends^29^, we set out to probe the sequence requirements for TnsB binding. Previous large-scale mutagenesis of VchCAST transposon ends revealed the importance of nucleotide identity at positions 6–9 and 13 of each TnsB-binding site^25^. Our structure shows that nucleotides at positions 6, 9, and 13 are within range to form base-specific hydrogen bonding with residues in the linker between DBD1 and DBD2 (**Figure 4A,B**). This linker region was previously proposed to contribute to V-K CAST transposon end recognition^26^, and it is strikingly similar to the AT-hook motif that contributes to Mos1 transposon end recognition^4^ (**Figure S13**). Superimposing the chains of this region from TnsB-LE1, TnsB-LE2 and TnsB-RE1 reveals a similar structure (**Figure 4C**), supporting the idea that these requirements apply to all TnsB binding sites^25^.

**Figure 4.**
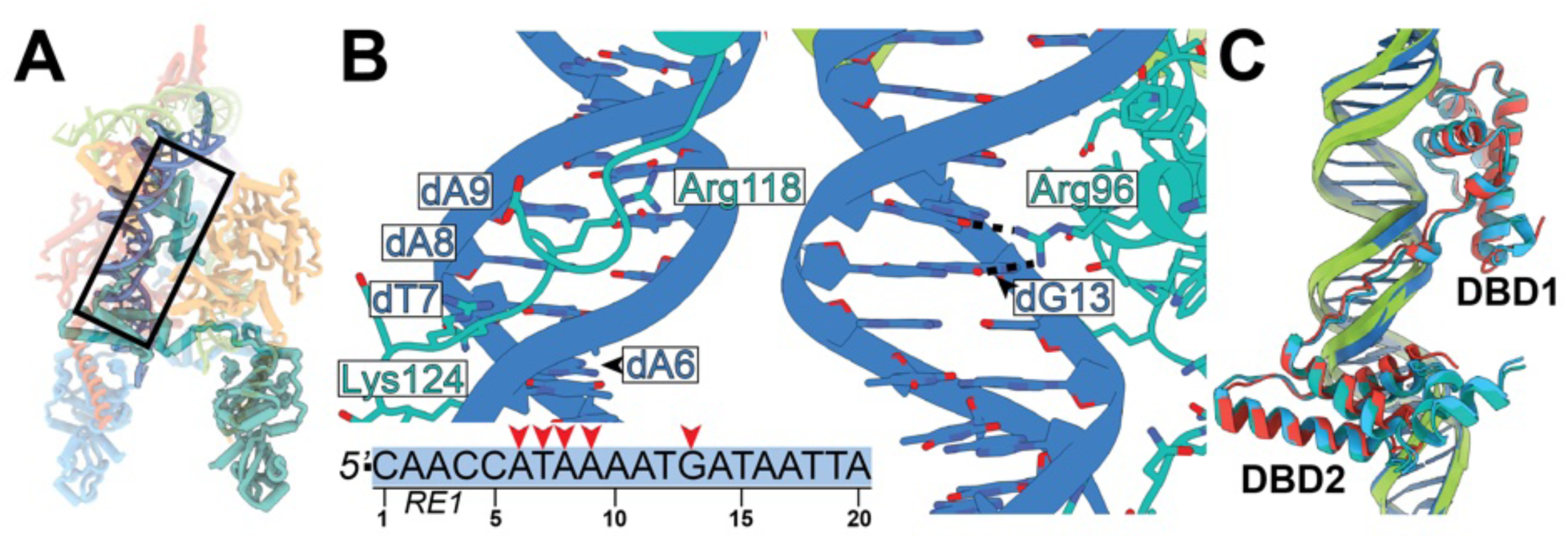
Base-specific interactions are associated with highly conserved positions within the DNA minor groove. Color scheme follows that of figure 1. **A.** Overview of the VchCAST complex structure. Black box marks regions shown in panel B. **B.** Linker between DBD1 and DBD2 (in cartoon and stick representation) and RE1 DNA. Bottom shows DNA sequence, with red arrows marking positions that cannot be altered without affecting activity. Protein residues that bind DNA are labeled. Hydrogen bonding interactions are indicated with black dashed lines. **C.** Superposition of the three DBD1-DBD2 regions from TnsB-LE1, TnsB-LE2 and TnsB-RE1 including linker regions and their bound DNA.

Earlier studies of the Tn7 transposase suggested that differences in binding affinity govern the sequential order of transposase binding during transposon-end synapsis^17,30^. We asked whether this also extends to the VchCAST system, and performed EMSA binding assays for each of the TnsB-binding sites (**Figure S14**). We determined that TnsB has similarly high affinity (Kd ∼5–10 nM) for all its binding sites (**Figure S14**), consistent with the fact that all DNA-binding domains are structured very similarly on their respective binding sites (**Figure 3C**). Therefore, we conclude that synaptic complex assembly is not driven by intrinsic differences in binding site affinity, at least for this particular CAST system.

## Discussion

Based on our findings, we propose a model in which IHF-induced DNA distortion drives formation of a cross-stabilizing TnsB interface that synapses the transposon ends, while V78 functions as a molecular “staple” that locks the complex into an integration-competent state(**Figure 5**). Cellular concentrations of IHF are high enough to ensure occupancy of its cognate binding site in the transposon left end^31,32^ (**Figure 5**). Through our structural and MD analysis, we propose that DNA distortions caused by IHF binding enable a transposase dimer to cooperatively bind LE1 and LE2. On the right end, the arrangement of the two terminal binding sites RE1 and RE2 would also promote cooperative binding of a TnsB dimer (**Figure 5**), as suggested by previous models based on Tn7^17^. The catalytic domain of the transposase bound to the first binding site on either end is likely flexible in this dimeric configuration, enabling the catalytic tetramer to form its characteristic ‘swapped’ configuration upon synapsis (**Figure 5**). In this synapsed arrangement, the proximity of the N-terminal DNA-binding domains (DBD2) forms cross-stabilizing interactions, which act as a ‘staple’ that secures the assembly and prevents reversible dissociation (**Figure 5**). In the absence of IHF (**Figure 5**), this tetrameric assembly can still form, but our MD analyses indicate that it would most likely be less stable and therefore less active than a left-right end pairing. Assembly of the catalytic core is a pre-requisite for donor DNA cleavage^33^, which is followed by genomic integration of cargo. Although the third transposase binding site on the left end is required for activity, these sites are not occupied in our structure. We speculate that this binding site may be transiently occupied earlier in the transposition process, or, alternatively, our sample preparation process prevents us from seeing full occupancy of all transposase binding sites (**Figure 5**).

**Figure 5.**
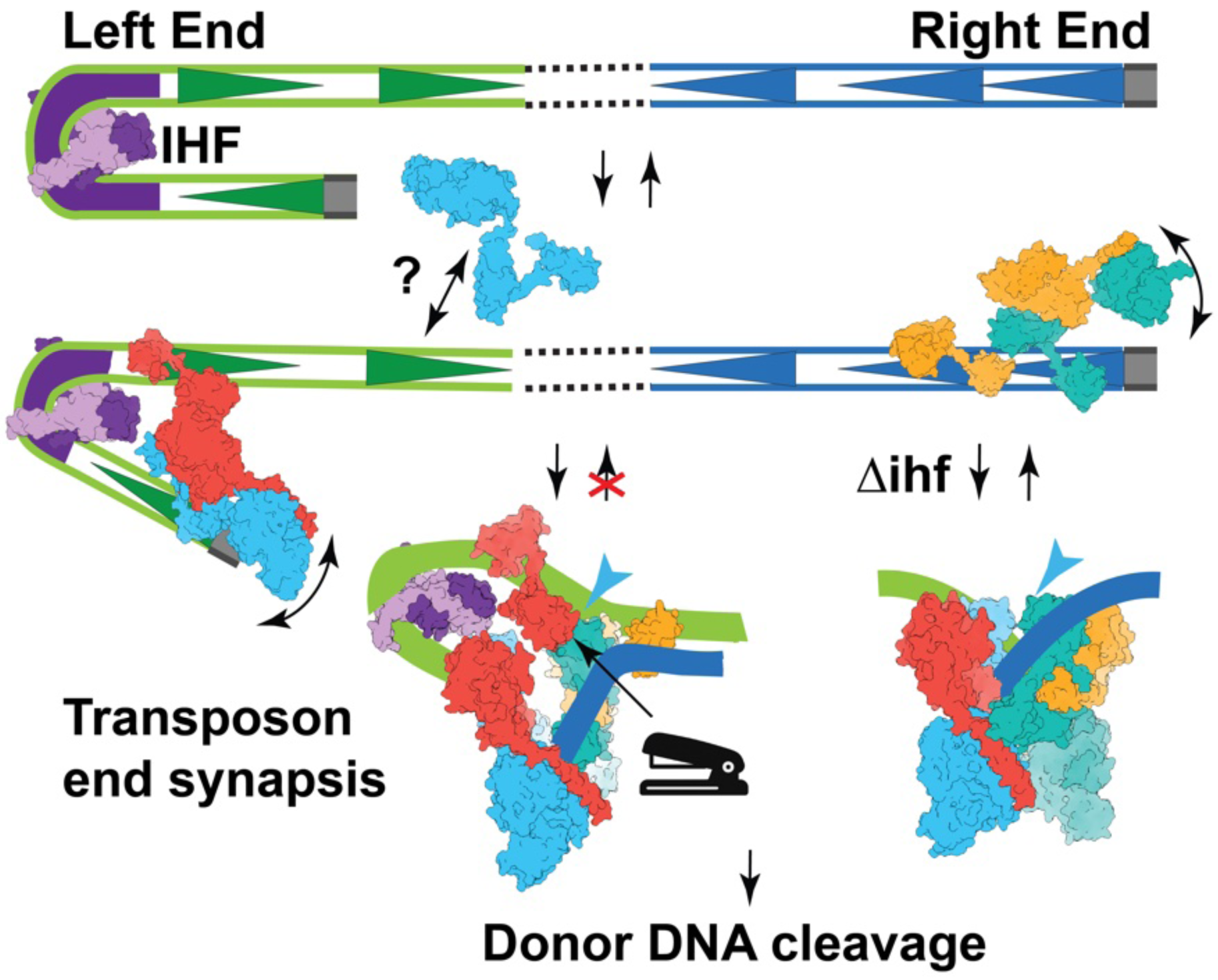
Model of transposon end pairing. Left (green) and right (blue) transposon ends are diagrammed, triangles represent TnsB-binding sites as shown in figure 1. Dotted line indicates intervening transposon DNA, host DNA is removed for clarity. IHF (purple) and TnsB protomers (light blue, red, yellow, and aqua) are at their respective binding sites. Single-headed arrows indicate the direction of the process. Straight double-headed arrows indicate binding/dissociation, and curved double-headed arrows indicate flexibility. Question mark indicates parts of the model that remain unsubstantiated. TnsB cartoons are drawn based on PDB ID 9Y7B (reported here) and 7PIK. Staple icon points to the interface formed between two TnsB subunits in the context of the paired end assembly. Light blue arrow indicates the location of the newly observed protein-protein interface formed in the presence of IHF. Two different paired end structures (the product of transposon end synapsis) are shown, the left diagrams the result of including IHF and the right the lack of IHF (indicated as τιIHF).

Transposons do not cross-mobilize the ends of even closely related homologs^29^, indicating transposon end recognition must be extremely stringent. Nevertheless, members of the I-F3 CAST family have similar transposase binding site configurations in their right and left ends^29,25^, suggesting that all I-F3 homologs adopt similar architectures. Furthermore, transposase structures^4,10,11,26,34^ consistently display very few sequence-specific contacts, suggesting that sequence specificity is not derived from direct interactions with the transposon end DNA. What, then, gives the transposase its exquisite specificity for its own ends?

Our observations address this paradox by suggesting that neither the structure of the transposase itself nor the specific interactions with any individual transposase binding site is sufficient for transposon end recognition. Instead, we propose that the geometry of the donor DNA itself, together with the placement of transposase binding sites, determines ‘allowable’ configurations of the assembly. Such constraints would be weakly dependent on the full transposon sequence and therefore could be subject to dramatic reconfiguration while still supporting robust levels of integration activity and specificity. The resulting DNA distortions may be induced through the coordinated action of the AT-hook like motifs, as proposed previously^35^. We conclude that specificity is weakly encoded in individual transposase binding sites but strongly encoded across all binding sites. Furthermore, the placement of binding sites for productive synapsis is subject to constraints imposed by the structure of DNA itself. This model suggests that: 1. transposon end identification could be improved by incorporating prior structural information and 2. transposon end sequences can be extensively redesigned while retaining high transposition activity, thereby increasing flexibility for cargo DNA and enabling precise in-frame edits. Unlike the catalytic core assembly, which remains relatively constant across the Tn7-like transposon superfamily, we anticipate that the architectural configuration we observe here is corresponds to one possible structure of potentially many possible synaptic architectures. Notably, while computational analyses of mutational changes prove useful here, we find that it is important to consider the impact of sequence changes at all possible TnsB-TnsB interfaces to fully understand sequence-function relationships. Thus, continued study of synaptic assemblies will likely yield deeper mechanistic insight into the fundamental rules governing transposon end synapsis and transposition.

## Material and Methods

The VchCAST *TnsB* gene (codon-optimized for *Escherichia coli*) was synthesized as a gift from Arzeda. The synthetic gene was cloned into the pARZ4 expression vector, which includes an N-terminal 6×His tag followed by an MBP (maltose-binding protein) tag and a TEV protease cleavage site. The plasmid was transformed into *E. coli* One Shot BL21 DE3 (Invitrogen) cells. Cultures were grown in 2×YT medium at 37 °C until an OD₆₀₀ of 0.4–0.6 was reached. Protein expression was induced with 0.25 mM isopropyl-β-D-thiogalactopyranoside (IPTG), and cultures were incubated overnight at 16 °C. Cells were harvested by centrifugation at 5,000 rpm for 20 min at 4 °C. Pellets were resuspended in lysis buffer containing: 50 mM Tris-HCl (pH 8.0), 500 mM NaCl, 1 mM EDTA, 0.1% NP-40, 1 mM DTT, and a protease inhibitor cocktail (Pierce, Thermo Scientific) at a ratio of 1 tablet per 30 mL buffer. Cells were lysed by sonication (7 cycles of 5 seconds on, 10 seconds off), and the lysate was clarified by centrifugation at 16,000 rpm for 30 min at 4 °C. Supernatant was applied to an amylose resin column (New England Biolabs) pre-equilibrated in wash buffer (50 mM Tris-HCl pH 8.0, 500 mM NaCl, 1 mM EDTA, and 1 mM DTT). The column was washed thoroughly, and the bound MBP-tagged protein was eluted with elution buffer containing 50 mM Tris-HCl (pH 8.0), 500 mM NaCl, 1 mM EDTA, 1 mM DTT, and 10 mM maltose.

Eluted fractions containing MBP-TnsB were pooled and diluted to 200 mM NaCl using Heparin dilution buffer (50 mM Tris-HCl pH 7.4, 50 mM NaCl, and 5% glycerol) and subsequently loaded onto a HiTrap™ Heparin HP column (Cytiva) on an ÄKTA Pure FPLC system. Protein was eluted using a linear salt gradient from 200 mM to 1,100 mM NaCl, created using Heparin low-salt buffer (50 mM Tris-HCl pH 7.4, 200 mM NaCl, 5% glycerol) and high-salt buffer (50 mM Tris-HCl pH 7.4, 1,100 mM NaCl, 5% glycerol). The eluted protein was further purified by size-exclusion chromatography using a Superdex 200 column (GE Healthcare) on the ÄKTA Pure FPLC system, equilibrated in gel filtration buffer containing 50 mM Tris-HCl (pH 7.4), 500 mM NaCl, and 5% glycerol. Protein purity was assessed by SDS-PAGE followed by Coomassie staining. Aliquots of the purified protein were flash-frozen in liquid nitrogen and stored at −80 °C.

### IHF purification

The pACYCDuet-1-ihfαβ co-expression plasmid was a gift from the Nuñez lab^36^. Both proteins were expressed by transforming the plasmid into One Shot BL21 DE3 (Invitrogen) cells. Cultures were grown in 2×YT medium at 37 °C until an OD₆₀₀ of 0.4– 0.6 was reached, followed by overnight induction at 16 °C with 0.25 mM IPTG. Cells were harvested by centrifugation at 5,000 rpm for 20 min at 4 °C and lysed by sonication (7 cycles of 5 seconds on/10 seconds off) in Ni-NTA lysis buffer containing 20 mM HEPES-NaOH (pH 7.4), 500 mM KCl, 0.1% Triton X-100, 2 mM DTT, 0.5 mM phenylmethylsulfonyl fluoride (PMSF), protease inhibitor cocktail, and 10% glycerol. The lysate was cleared by centrifugation, and the soluble fraction was loaded onto a HisTrap™ HP column (Cytiva) using an ÄKTA Pure FPLC system. His-tagged proteins were eluted with an imidazole gradient generated using Ni-NTA buffer A (20 mM HEPES-NaOH pH 7.4, 500 mM KCl, 10 mM imidazole, 5% glycerol, 1 mM DTT) and Ni-NTA buffer B (same as buffer A but with 500 mM imidazole). The eluted protein was digested with TEV protease overnight at 4 °C to remove the His tag. The IHF heterodimer was further purified using a HiTrap™ Heparin HP column (Cytiva) with a linear KCl gradient from 250 mM to 1,000 mM. Peak fractions were pooled and concentrated using a 3 kDa molecular weight cutoff concentrator and subjected to size-exclusion chromatography on a Superdex 75 16/60 column (GE Healthcare), equilibrated in gel filtration buffer containing 20 mM HEPES-NaOH (pH 7.4), 500 mM KCl, 5% glycerol, and 1 mM DTT. The purified IHF heterodimer was analyzed by SDS-PAGE followed by Coomassie staining. Aliquots were flash-frozen in liquid nitrogen and stored at −80 °C.

### Preparation of DNA substrate

DNA substrate was designed similarly to previously reported transpososome substrates used for structural studies^37^. Briefly, an asymmetric DNA substrate that mimics the VchCAST RNA-guided transposition product was made comprising two DNA substrates: (1) one containing 105 base pairs of the native transposon left-end sequence, a target site duplication, and a flanking genomic region; and (2) a separate substrate containing 47 base pairs of the transposon right-end sequence, a target site duplication, a flanking target region, and a desthiobiotinylated LUEGO sequence for complex isolation. The left-end substrate is composed of three single-stranded DNA oligos, while the right-end substrate is composed of four. Prior to annealing, oligos were purified by high-performance liquid chromatography (HPLC) using an Agilent 1260 Infinity II system equipped with an AdvanceBio Oligonucleotide column (Agilent Technologies) to ensure incorporation of full-length oligos into the substrate. For substrate preparation, oligos (IDT) were mixed in an equimolar ratio supplemented with a 10X concentrated annealing buffer for a final buffer composition of 10 mM Tris pH 7.5, 50 mM NaCl, and 1 mM EDTA.

The mixture was then heated to 95 °C for 5 min and cooled down to 50 °C at the rate of 1 °C per min using a thermal cycler (BioRad).

### Synaptic Complex Reconstitution

Two sample reconstitutions were performed to form the TnsB synaptic end complex. The transpososome substrate was mixed with TnsB and IHF in a 1:10:16 stoichiometric ratio. The final buffer composition of the reconstitution was 26 mM HEPES, pH 7.5; 5 mM Tris-HCl, pH 7.5; 20 mM KCl; 750 mM NaCl; 0.2 mM MgCl₂; 15 mM MgOAc; 2% glycerol; 1 mM ATP; 1 mM DTT; and 0.1% Tween-20. The sample was dialyzed overnight at 4°C into a dialysis buffer with identical composition, except for the concentration of NaCl, which was lowered to 150 mM, hereafter referred to as low-salt buffer. Streptavidin-affinity beads (Streptavidin Mag Sepharose magnetic beads, Cytiva) were washed three times with 500 µL of low-salt buffer before being incubated with the post-dialyzed sample for one hour at room temperature under rotation. The complex was isolated from DNA-unbound proteins by washing the beads three times with 500 µL of low-salt buffer, followed by elution using two rounds of 50 µL elution buffer containing low-salt buffer supplemented with 10 mM biotin. Elution buffer was incubated with the beads under rotation for 30 min for each round. Elutions were analyzed via SDS-PAGE gel to confirm the presence of TnsB and IHF.

### Cryo-EM Sample Preparation and Imaging

The initial synaptic end complex reconstitution samples were prepared for cryo-EM by diluting the pull-down elution two-fold, four-fold, or by using the undiluted elution. Custom graphene grids were prepared following a previously described protocol^38,39^. To render the grids hydrophilic, graphene grids were plasma-cleaned in the Solarus II Plasma Cleaner (Gatan) under Ar/O2 at 10 W for 10 seconds with the graphene coated surface facing upwards in the chamber^40^. Four microliters of the transpososome complex sample were loaded onto the carbon side of a graphene-coated grid and incubated for 60 seconds in the Mark IV Vitrobot chamber (ThermoFisher), which was set to 4°C and 100% humidity. Each grid was blotted for 3.5 seconds with a blot force of –7, then plunged into liquid ethane cooled with liquid nitrogen. The second synaptic end complex reconstitution elutions were frozen on UltrAuFoil grids (1.2/1.3, 300 mesh, QuantiFoil). UltrAuFoil grids were plasma-cleaned under Ar/O2 at 10 W for 30 seconds in the Solarus II Plasma Cleaner. Four microliters of undiluted elution samples were loaded onto the carbon side of an UltrAuFoil grid in a Mark IV Vitrobot chamber at 4°C and 100% humidity. Each grid was blotted for 3.5 seconds at a blot force of –7, then plunged into liquid ethane cooled with liquid nitrogen.

### Cryo-EM imaging and image processing of the transposase complex

Vitrified samples of the transposase complex were first screened on a Talos Arctica (Thermo Fisher Scientific) equipped with a Gatan K3 direct electron detector and a BioQuantum energy filter, prior to large-scale data collection. Sample grids were imaged at 200 kV, with an intended defocus range of −2.5 to −1.0 μm and a magnification of 130,000x in electron counting mode (0.63 Å per pixel). Movies were collected with a total dose of 60 e-per Å^2^. A total of 22,812 micrographs were collected using EPU.

Downstream processing was performed in cryoSPARC 4.7.0^41^. Movies were motion-corrected and summed using Patch motion correction in cryoSPARC. The contrast transfer function (CTF) was estimated using PatchCTF in cryoSPARC. A total of 8,000 micrographs were initially processed. Particle picking was performed using blob picker, followed by extraction and 2-D classification. Particles from representative complex classes were used for training the Topaz model to pick from the entire dataset. Particles were extracted with a 512-pixel box and subjected to multiple cycles of refinement to remove junk particles. Heterogenous refinement was initially performed against two classes, a STC reference model and a junk class. The resulting best class was used for local refinement focused on three regions: the IHF-bound site, the catalytic core, and distal end DNA regions. The composite map was generated from these local refinement maps using UCSF ChimeraX v1.8 command ‘vop maximum’ after aligning the local reconstructions^42,43^. The resolution was estimated using the gold-standard method. Local resolution was estimated using cryoSPARC. CryoSPARC 3D Flex tools were used to analyze sample heterogeneity, with the output mode set to 41 frames^44^. The resulting series was visualized in UCSF ChimeraX v1.8 using the vseries command to display all frames as a movie. The results are presented in Supplementary Video 1.

### Model Building and Refinement

The model was built in a multi-step process. Individual TnsB domains from the AlphaFold3 model were first rigid body docked into locally refined maps^27^. Further adjustments within domains were made using Coot rigid body and real space refine utilities^45^. TnsB domains were then rigid body docked into a composite map constructed from three locally refined maps focused on three separate regions; the catalytic core, the IHF bound site, and the distal left end segment. The DNA model was largely constructed from combining DNA associated with the catalytic core from an AlphaFold3 complex prediction^27^, bent IHF-bound DNA (AlphaFold3 prediction of DNA substrate complex with IHF) and the distal left-end DNA sequence (fragments from PDB entry 7PIK). TnsB domain linkers were then taken from the AlphaFold3 model and placed into the composite map using rigid body followed by real space refine in Coot^46^. The TnsB-DNA model was then refined with Phenix using a low-resolution protocol, which included DNA base-pair restraints^47^. IHFα and IHFβ from the IHF/DNA complex crystal structure (1IHF) was then rigid body docked into density individually. Within each protomer, the β-hairpin portion was rigid body fit separately to better fit the density. Minor adjustments to the model were made to alleviate clashes with the DNA, mostly at the proline intercalation site. The entire model was then refined and B-factors calculated using Phenix (refer to Table1 for refinement statistics).

### Mate-in *in vivo* transposition assay

Mate-in assays were performed using two E. coli strains: a BL21 strain (New England Biolabs) containing two helper vectors, and a PO603 strain carrying a donor plasmid with a mini-element. The helper vectors in the BL21 strain encode Tn6677 TnsA-TnsB-TnsC (Addgene #130633, modified to confer ampicillin resistance) and TniQ-Cas8-Cas7-Cas6-sgRNA (Addgene #130635). The second helper vector is cloned to contain a LacZ spacer within the sgRNA to target a genomic region in the BL21 strain. The mini-element within the donor plasmid contains a kanamycin resistance gene flanked by canonical VchCAST transposon ends, and the plasmid contains an R6K origin to prevent replication within the BL21 strain. The donor cell strain (PO603), a gift from Joseph E. Peters (unpublished), contains an R6K origin of replication and an origin of transfer. TnsB mutation vectors (V78A, V78D, V78K, B78F, and catalytic mutant D308A) were generated via site-directed mutagenesis (New England Biolabs) using the parental TnsABC helper vector. Each strain was grown in 50 mL Luria–Bertani (LB) medium supplemented with 0.1 mM IPTG and appropriate antibiotics (100 μg ml^−1^ carbenicillin, 25 μg ml^−1^ spectinomycin, 50 μg ml^−1^ kanamycin) to an optical density (OD) of 0.6 in a 37°C shaking incubator. Cultures were then centrifuged at 4,000 rcf for 10 min to pellet the cells. The supernatant was removed, and pellets were resuspended in 20 mL LB containing 0.1 mM IPTG. This washing step was repeated once more to remove residual antibiotics. After the second wash, cells were resuspended in 500 µL LB with 0.1 mM IPTG. 10 uL of each resuspension was diluted 50-fold with LB for OD_600_ measurements using the UV5Bio Spectrophotometer (Mettler Toledo) to determine cell density. The measurements were used to mix recipient and donor resuspensions in a 1:5 ratio, with a final volume of 150 μL. The mixtures were vortexed for 10 seconds, centrifuged at 15,000 rcf for 1 min, and incubated at 30°C for 24 hours. Following incubation, cells were plated at 1x and 10x dilutions onto LB-agar plates containing 100 μg ml^−1^ carbenicillin, 25 μg ml^−1^ spectinomycin for cells, and 50 μg ml^−1^ kanamycin. Antibiotic-resistant colonies were counted after overnight incubation.

### Molecular Dynamics Simulations

Molecular dynamics simulations were based on the cryo-EM structure of full TnsB-IHF-DNA ternary complex from *Vibrio Cholerae* (PDB ID: 9Y7B). Missing protein segments were modelled using AlphaFold3^27^ and Swiss-Model^48^. To obtain the TnsB-DNA complex without IHF, the IHF heterodimer was removed from the full structure. In addition, two simplified systems containing only the LE-DNA segment, in the presence or absence of IHF, were constructed to assess the specific contribution of IHF to DNA architecture. Each system was solvated in explicit water, and counter ions were added to neutralize the total charge yielding ∼627 K atoms for the full ternary complex and ∼238 K atoms for the simplified systems.

All simulations employed a protocol optimized for RNA/DNA nucleases. The Amber ff19SB force field^49^ was used for proteins, together with the ff99bsc1 corrections for DNA^50^ and the ff99bsc0+χOL3 parameters for RNA^51^. Water molecules were described using the TIP3P model^52^. An integration time step of 2 fs was employed and the bonds involving hydrogen atoms were constrained using the SHAKE algorithm^53^. Temperature was maintained at 300 K using Langevin dynamics^54^ and pressure was regulated at 1 atm using a Berendsen barostat^55^ with a 2 ps relaxation time. Initial energy minimization was performed to relax the solvent and counterions while holding protein and DNA atoms fixed with a 300 kcal·mol^-^¹·Å^-^² harmonic restraint. All the systems were subsequently equilibrated through a staged heating protocol, gradually raising the temperature to 300 K under positional restraints that were progressively released. Finally, production simulations were carried out in the NVT ensemble, with three independent replicates of 1 μs for each run. All trajectories were generated using the GPU-accelerated implementation in AMBER20^56^.

Distance analysis considered the center of mass of the V78 residue from TnsB-LE2 and TnsB-RE1. Contact analyses were performed for TnsB-LE2 versus TnsB-RE1 and for TnsB-LE2 versus TnsB-LE1. A contact was considered when the distance between two heavy atoms among the regions of interest was < 5 Å. Kernel density estimates were generated using the kdeplot function. The 3D representations of the complex were rendered using VMD 1.9.3^57^.

## Data availability

Cryo-EM composite map and model for TnsB-IHF-DNA ternary structure were deposited in the EMDB and PDB, respectively, with accession codes EMD-72650 and PDB 9Y7B. All cryo-EM reconstructions used to generate the composite map are available through the EMDB with accession codes: EMD-72640 (consensus map), EMD-72641 (local refinement map of IHF-bound DNA region), EMD-72647 (local refinement map of distal DNA region), and EMD-72648 (local refinement map of catalytic core).

## Contributions

E.H.K. led project conceptualization, data analysis, and supervised the project. S.F. purified all proteins, V.H.T. and S.F carried out complex reconstitutions for cryo-EM sample preparation. Y.S. provided custom-made graphene grids for cryo-EM sample preparation. V.H.T. prepared cryo-EM samples, and collected and processed cryo-EM data. D.J.M. and V.H.T. built and refined atomic models. V.H.T. carried out functional assays. C.P. and A.F. performed molecular dynamics simulations. C.P., A.F., and G.P. analyzed the simulations data. E.H.K. wrote the initial manuscript draft, all authors edited and reviewed the manuscript.

## Acknowledgements

We gratefully acknowledge the valuable discussions and comments from Ines Chen, Shubham Dubey, and the rest of the Kellogg lab. We sincerely acknowledge the St. Jude Protein Purification Center, especially Richard Heath and Rosario Mosca for their valuable support and assistance with protein purification. We also gratefully acknowledge the Nuñez lab and Arzeda for providing constructs used for purification. This work was performed at the St. Jude Cryo-EM Center, which is generously supported by the St. Jude Children’s Research Hospital. This work was supported by NIH NIGMS 5R01GM144566 (E.H.K.), the Pew scholars program (E.H.K.), the Cystic Fibrosis Foundation (E.H.K.), NIH NIGMS F31GM151863 (V.H.T.), NIH (Grant No. R01GM141329 to G.P.), NSF (Grant No. CHE-2144823 to G.P.), and ALSAC, the fundraising and awareness organization of St. Jude Children’s Research Hospital (E.H.K.). GP acknowledges support by the Alfred P. Sloan Foundation (Grant No. FG-2023-20431) and the Camille and Henry Dreyfus Foundation (Grant No. TC-24-063). The computational studies performed here were carried out using the Pittsburgh Supercomputer Center through allocation BIO230007 from the Advanced Cyberinfrastructure Coordination Ecosystem: Services & Support (ACCESS) program, which is supported by NSF support grants #2138259, #2138286, #2138307, #2137603, and #2138296.

## Supplemental Figures

**Supplemental Figure 1.**
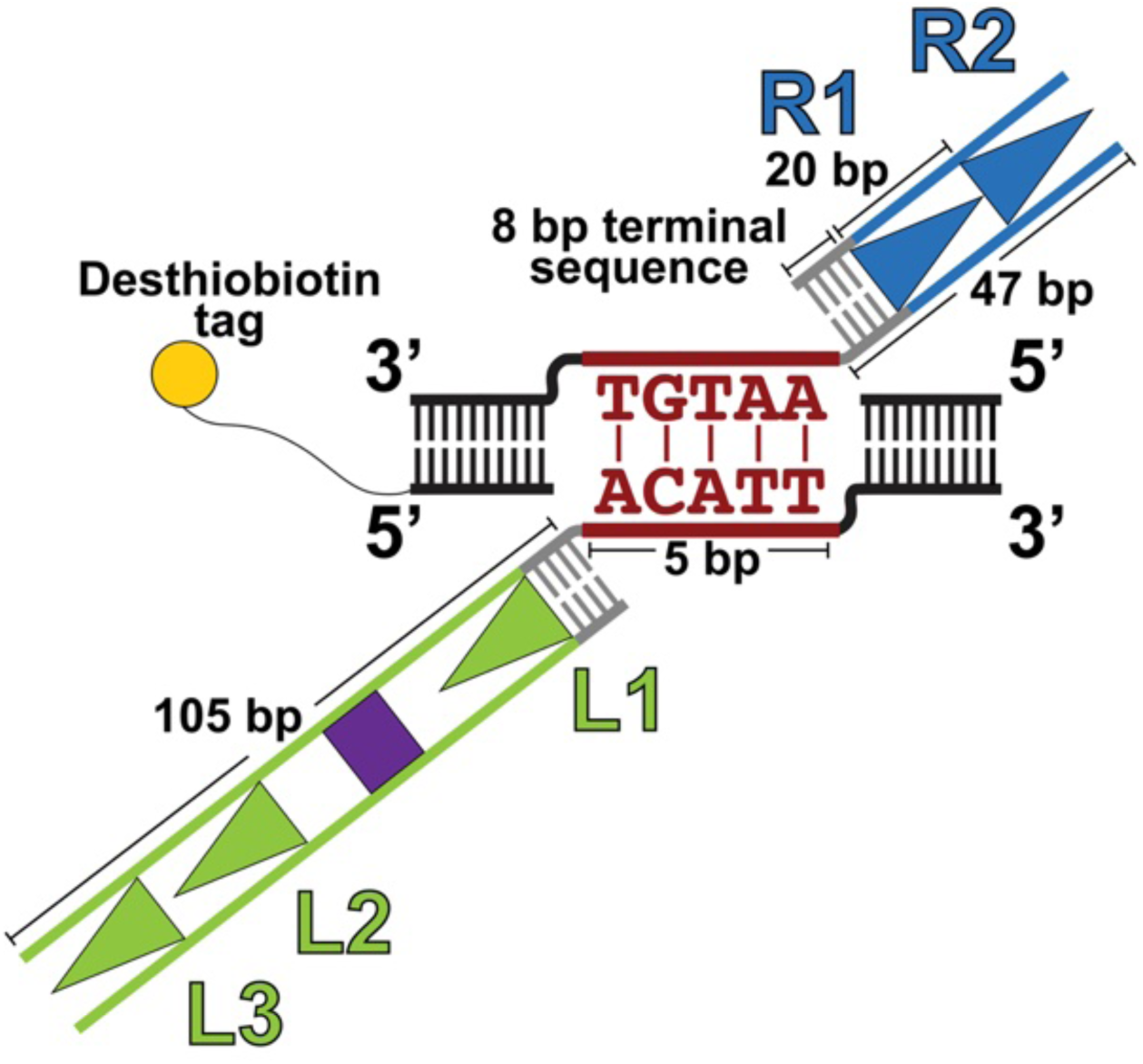
Schematic of the designed asymmetric strand-transfer DNA substrate, representing the Shapiro intermediate product of the VchCAST strand-transfer reaction. The 105-bp segment of the transposon left end (green) contains three TnsB binding sites (L1–L3; each site is 20 nucleotides, indicated by triangles) and an IHF-binding site (purple). The 47-bp segment of the transposon right end (blue) includes two TnsB binding sites (R1–R2). Each end contains an 8-bp terminal sequence (gray), while the target DNA is shown in black. A desthiobiotin tag (yellow circle) is attached to the target DNA to facilitate affinity pulldown during sample reconstitution.

**Supplemental Figure 2.**
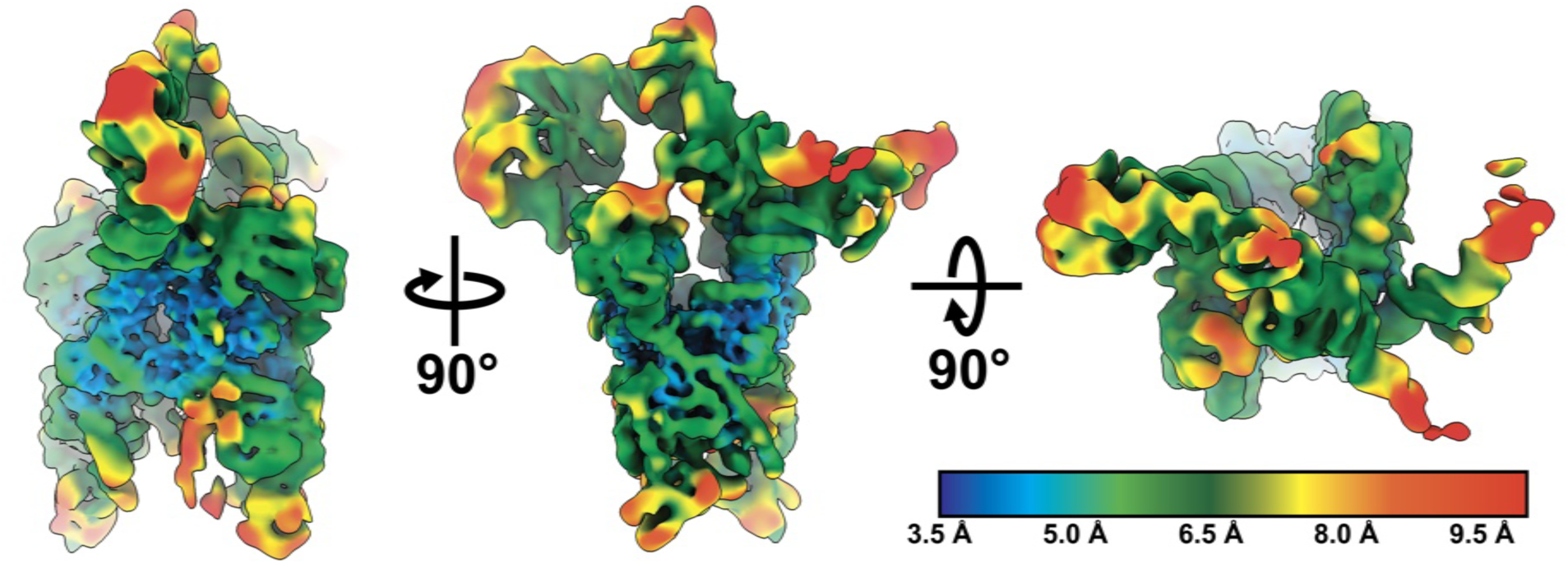
Local resolution map of consensus map. Map voxels are colored according to local resolution estimates; color legend is on bottom right of figure. Three different views of the locally filtered consensus map are shown, labeled by rotations required to relate the different views. Map shown corresponds to the locally filtered consensus map, displayed at threshold = 0.0936. Local resolution was assessed using the local resolution estimation tool in CryoSPARC, applying a Fourier shell correlation (FSC) threshold of 0.5.

**Supplemental Figure 3.**
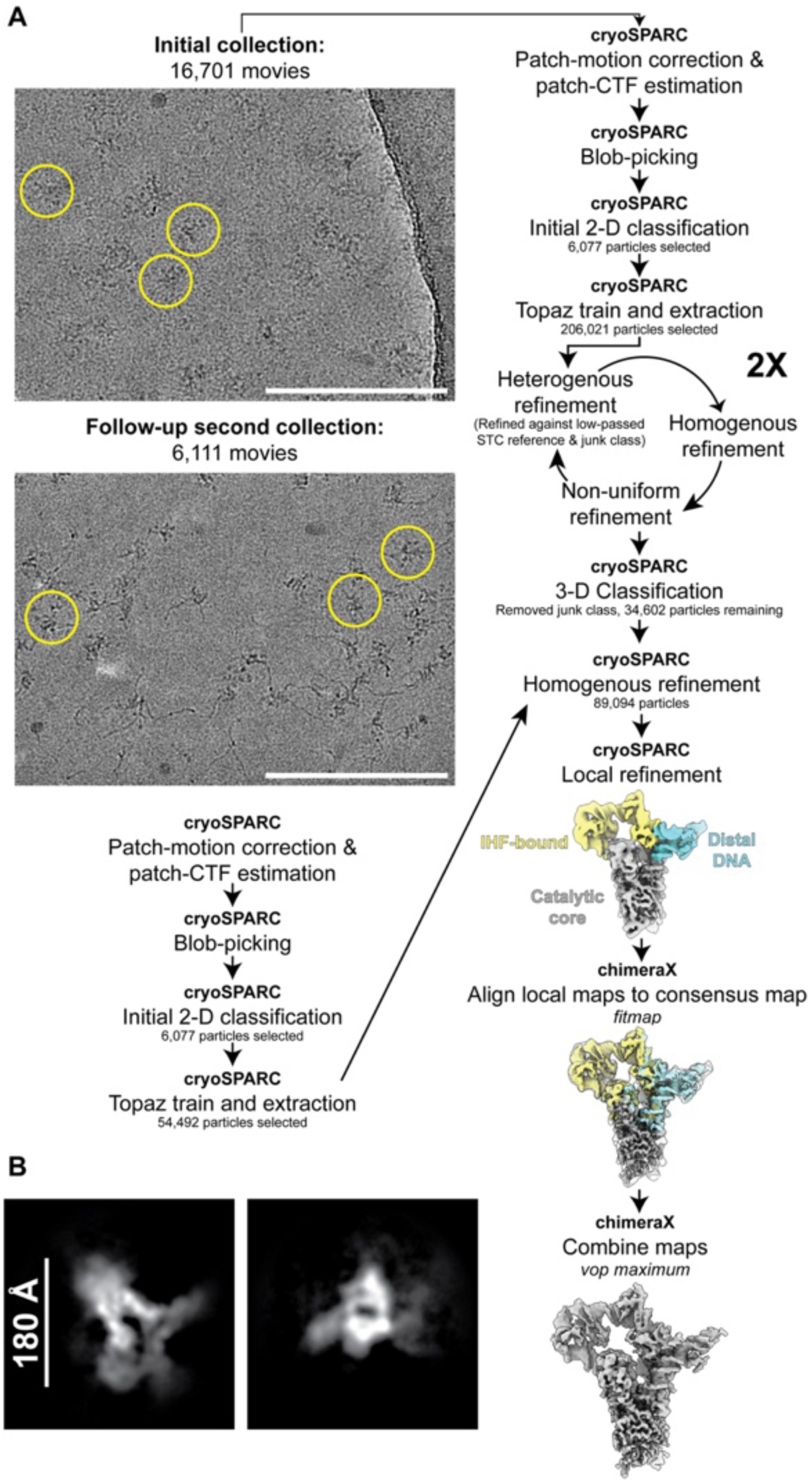
Image processing workflow to produce composite map. Representative micrographs are shown from each data collection session, with yellow circles highlighting examples of particles. White scale bar represents 150 nm. Image processing workflow from pre-processing (top) to final composite map reconstruction (at bottom). The output of three separate local refinement jobs focused on different regions: IHF-bound region, catalytic core, and distal end DNA. Locally refined maps are shown below in different colors: yellow, grey, and light blue respectively. **B.** Reference-free 2D classes from collected dataset. Scale bar represents 180 Å.

**Supplemental Figure 4.**
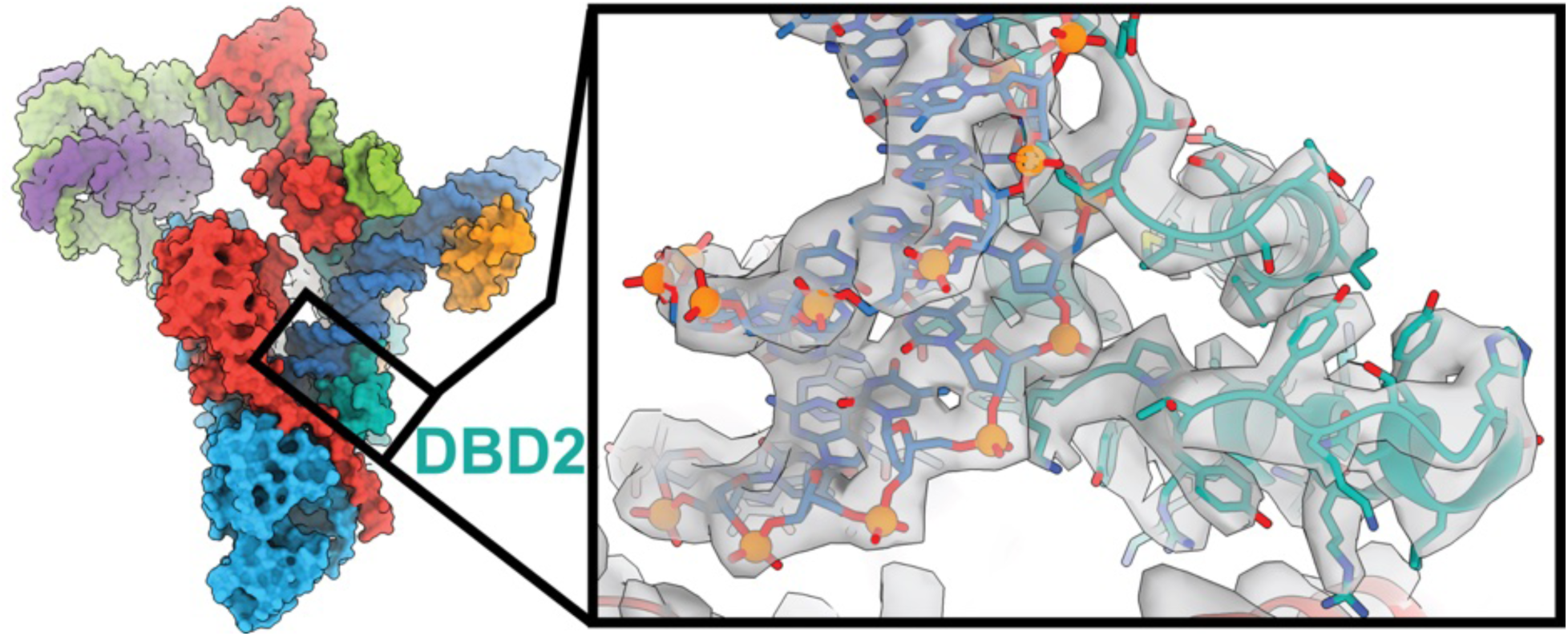
Highest resolution structural features observed in the catalytic core. Left: structure overview provided as a reference for the reader, boxed region indicates location of close-up, along with a label for the domain being visualized. Right: Close-up view of the atomic model, shown in sticks, of DBD2 (teal) associated with RE1 (blue) superimposed with the locally-refined map of the catalytic core (gray surface). Atoms are colored according to element: oxygen is orange, nitrogen is blue, phosphates are orange. The map threshold (threshold = 0.622) was chosen to highlight the separation between DNA bases.

**Supplemental Figure 5.**
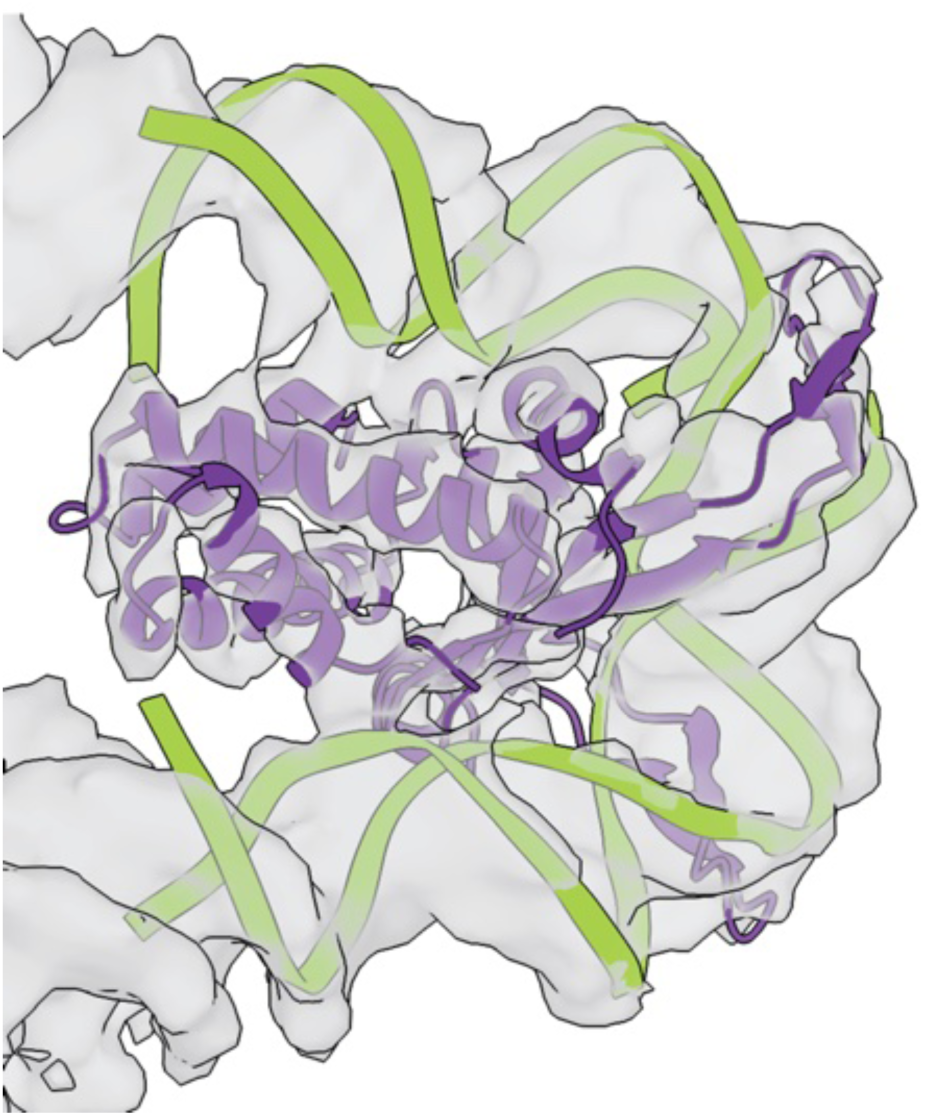
The crystal structure of previously determined IHF structure docked into the composite map. The co-crystal structure of *E. coli* IHF-DNA (PDB 1IHF) is rigid body docked into the VchCAST locally-refined IHF-bound region composite map (transparent gray surface, threshold = 0.44), showcasing the 180° DNA bend generated by the host factor. Protein is shown in purple cartoon, DNA backbone displayed in green ribbon.

**Supplemental Figure 6.**
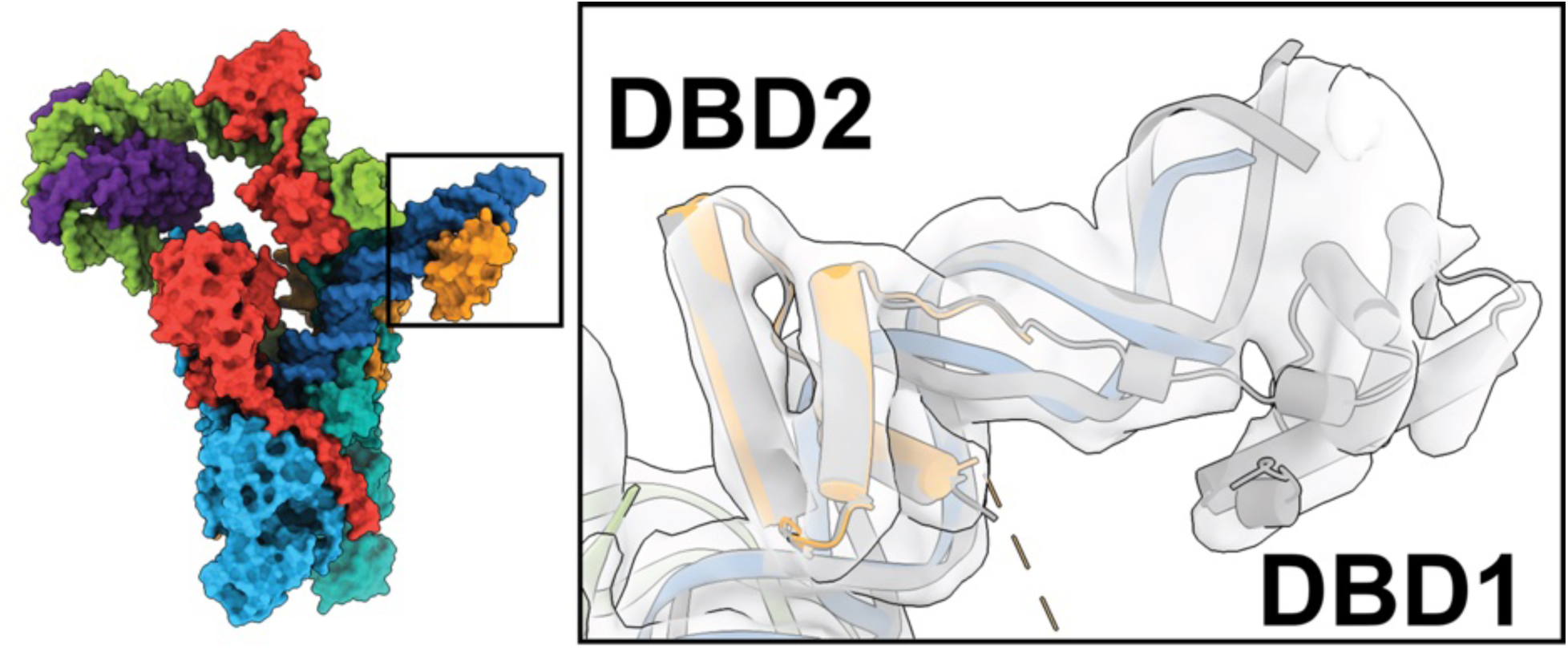
Expected location of DBD1 on RE2. Left: structure overview shown as a reference for the reader, boxed region indicates location of the close-up. Right: Locally filtered consensus cryo-EM map shown in transparent gray (threshold = 0.087). Although the DBD1 domain of TnsB-RE2 was not built due to weak density, superposition with the same region of TnsB-LE1 (cartoon representative, colored gray, DBD1 and DBD2 labeled) shows its expected location.

**Supplemental Figure 7.**
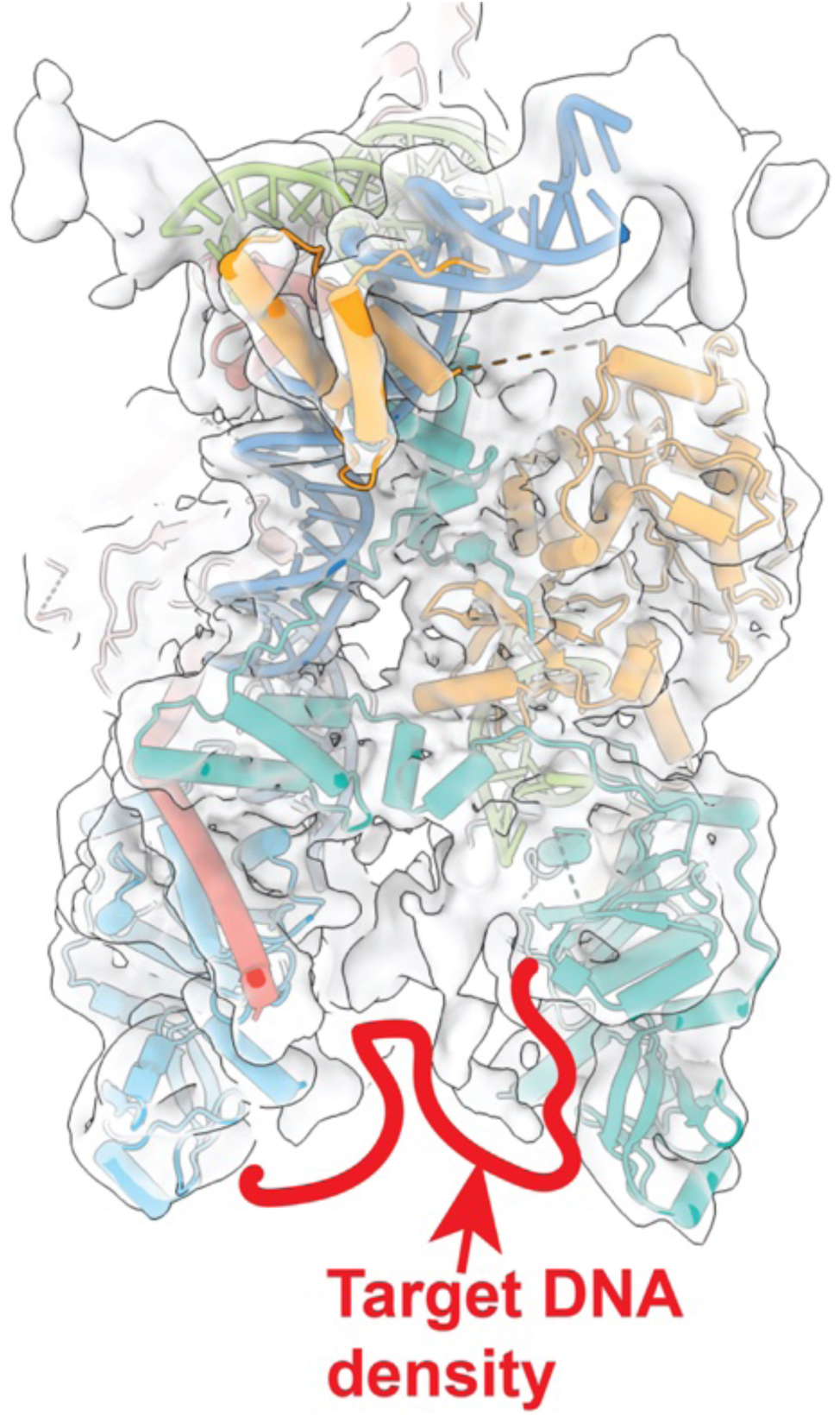
Cryo-EM map exhibits weak target DNA density. Locally filtered consensus map displayed at 0.082 threshold, in the same viewing direction as shown in Figure 2A. Map shown in transparent gray surface. Atomic model, shown in cartoon is docked into the map, colors are the same as throughout the manuscript. Target DNA density is partially outlined by the red line to facilitate visualization.

**Supplemental Figure 8.**
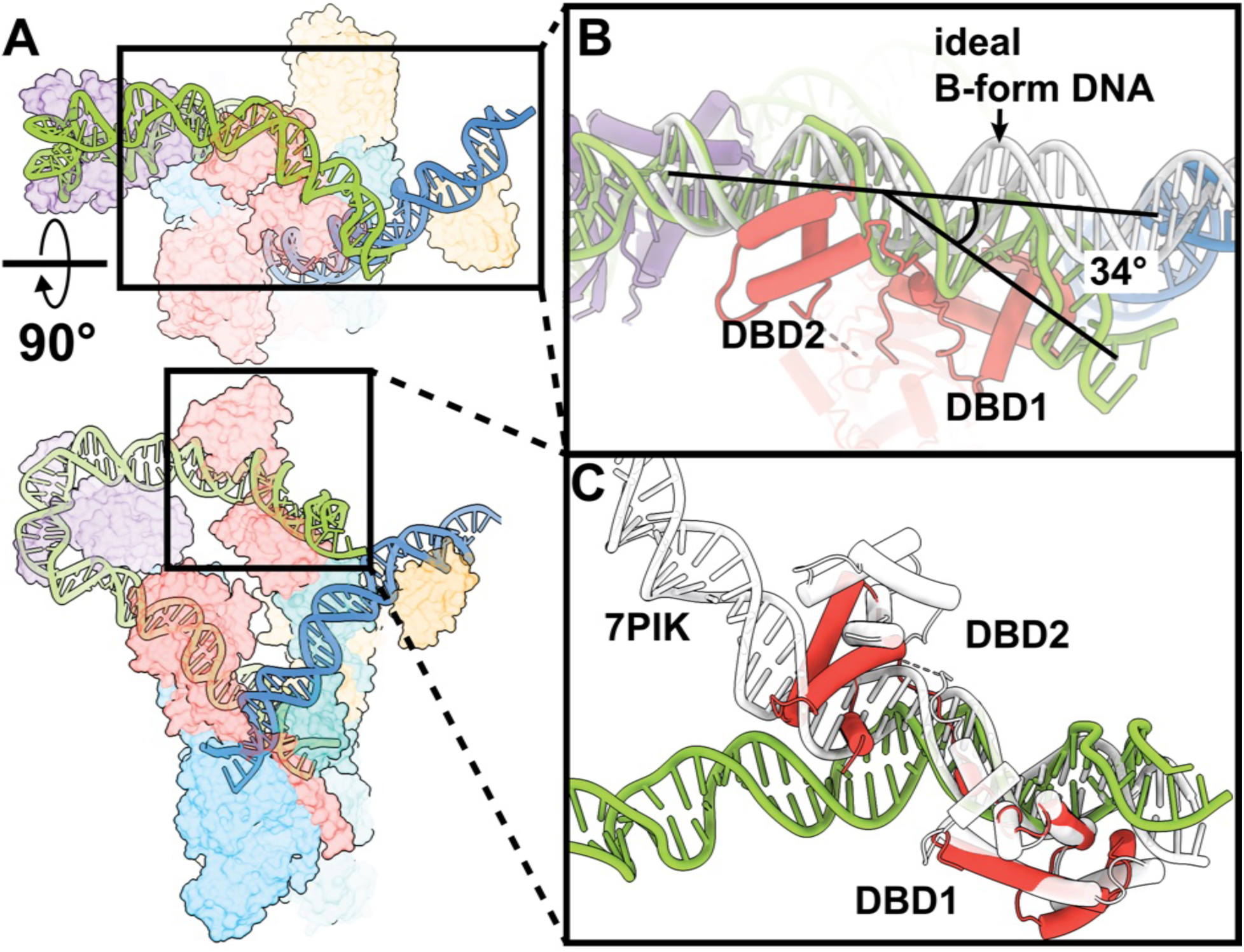
Local DNA distortions observed and compared to previously determined structures. **A.** Transparent surface representation of the atomic structure shown in two orthogonal views. Color scheme follows that of figure 1. DNA structure is shown in cartoon to facilitate visualization. Callouts (boxes) indicate the portions of the structure shown in greater detail in panels B and C. Rotation axis/angle indicates the rotation in view going from the bottom panel to the top panel. **B.** Cartoon representation of the atomic structure superimposed onto a model of ideal B-form DNA (white, ribbon). The left end is colored light green, and DNA-binding domains (DBD1 and DBD2) are labeled. The bending angle between ideal B-form DNA and the left end is shown (black lines) on top of the displayed structures. **C.** Previously determined Tn7 TnsB structure (PDB: 7PIK, white cartoon) is superimposed onto DBD1 of TnsB-LE2 (colored cartoon). DBD1 and DBD2 are labeled.

**Supplemental Figure 9.**
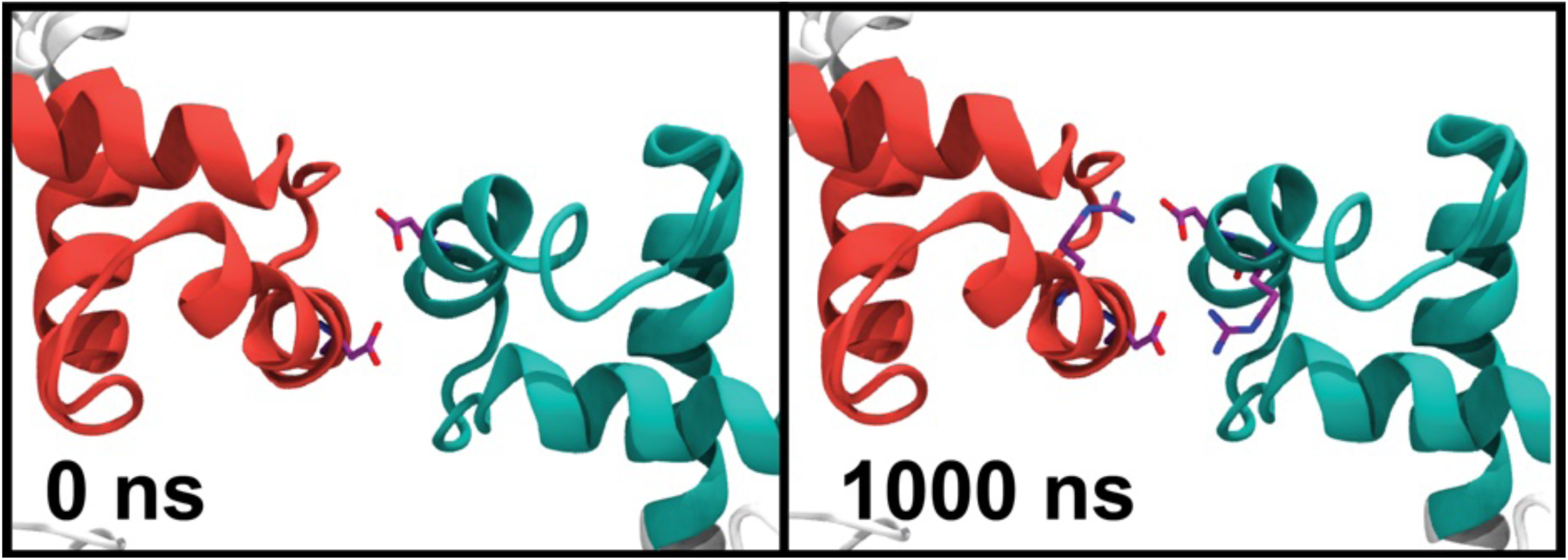
Molecular dynamics (MD) simulation snapshots of the V78D mutant showing compensatory interaction of V78D residues with neighboring Arginine residues (R38) at the TnsB-LE2, TnsB-RE1 interface, resulting in lack of separation. Timepoint in nanoseconds is indicated at the bottom left of each panel.

**Figure S10:**
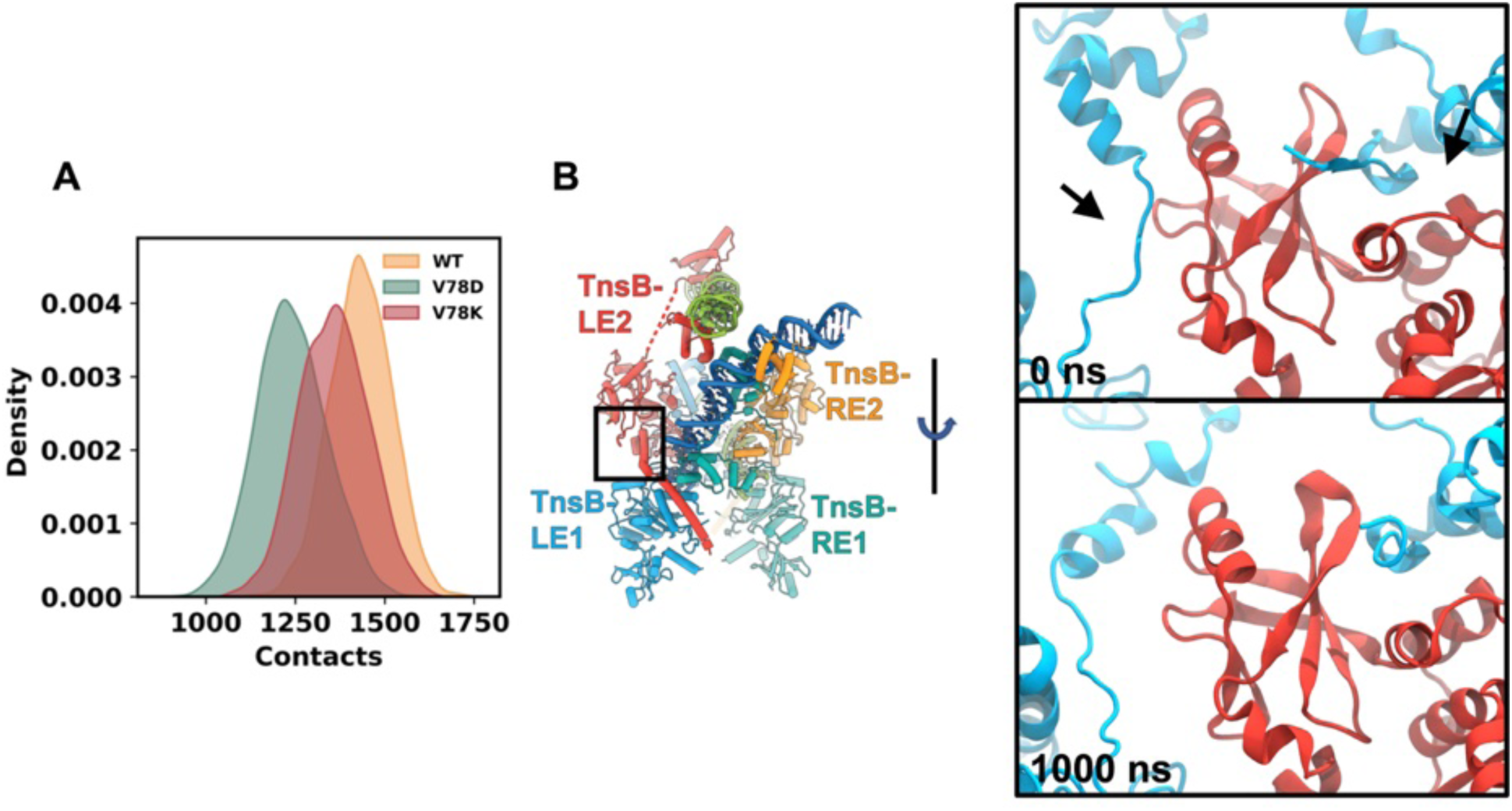
**A.** Density plots showing number of contacts between of LE2–LE1 inter-subunit from MD simulations of WT and mutant system. **B.** left: overview of interface being analyzed in panel A. right: MD snapshots of the V78D mutant, arrows highlight the regions at TnsB-LE2, TnsB-RE1 interface that contributed to reduction in contacts.

**Supplemental Figure S11.**
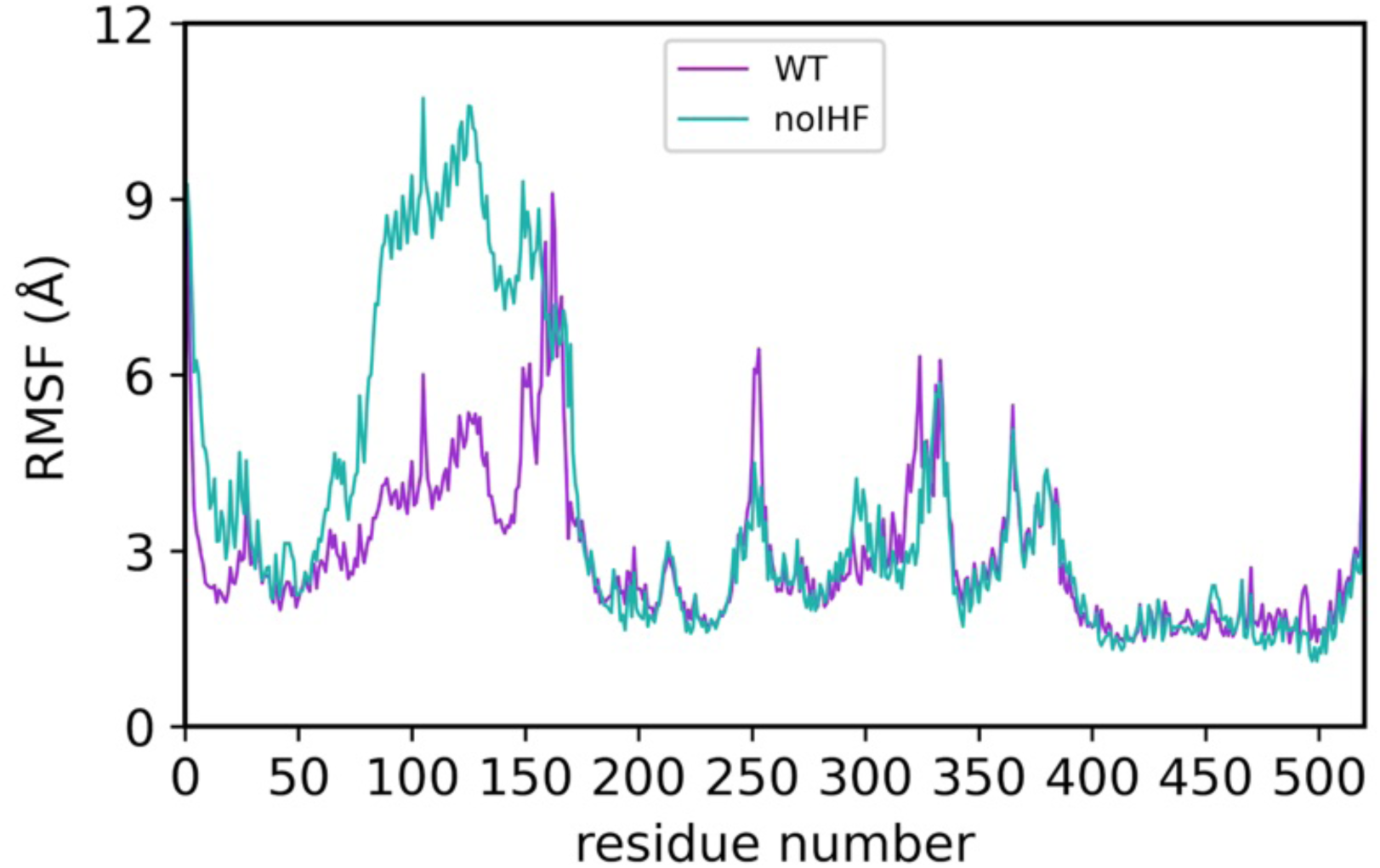
RMSF on a per-residue basis for TnsB-LE2. Residue number plotted on the x-axis, root-mean-square-fluctuation (RMSF) on the y-axis. WT include the full assembly with IHF bound (pink), noIHF corresponds to the the assembly lacking IHF (green).

**Supplemental Figure 12.**
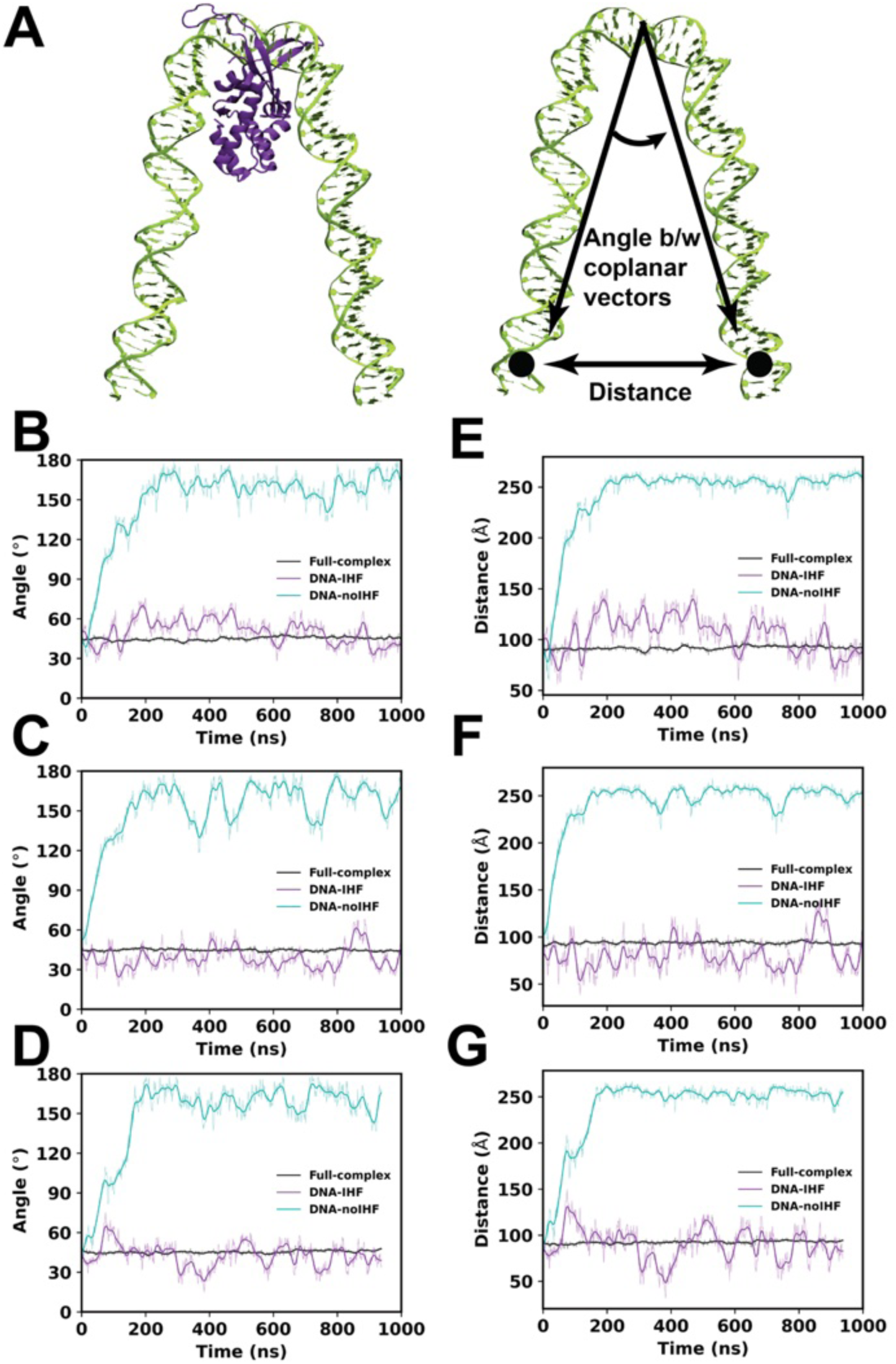
IHF stabilizes the LE-DNA conformation. Atomic model of the LE-DNA in presence of IHF (left panel) and in absence of IHF (right panel). A schematic illustration showing two coplanar vectors defined along the DNA. The angle between the vectors (curved arrow) captures their relative orientation, while the linear distance between their base points (black circles) represents the end-to-end distance. **B–D,** Time evolution of the vector angle for three independent replicas of LE-DNA simulated in the presence and absence of IHF. **E–G,** Corresponding end-to-end distances for the same replicas, illustrating the effect of IHF binding on LE-DNA conformation.

**Supplemental Figure 13.**
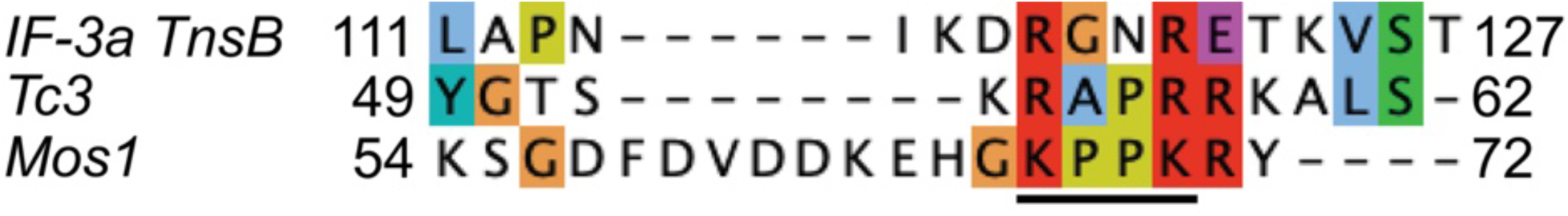
‘AT-hook’-like motif and DBD1-DBD2 linker sequence alignment. Sequence alignment of homologous segments from related transposons: Tc3 and Mos1. ‘AT-hook’-like motif is underlined. Residues are colored according to their biophysical properties and conservation, with the bounding residue numbers shown at the beginning and end of the alignment.

**Supplemental Figure 14.**
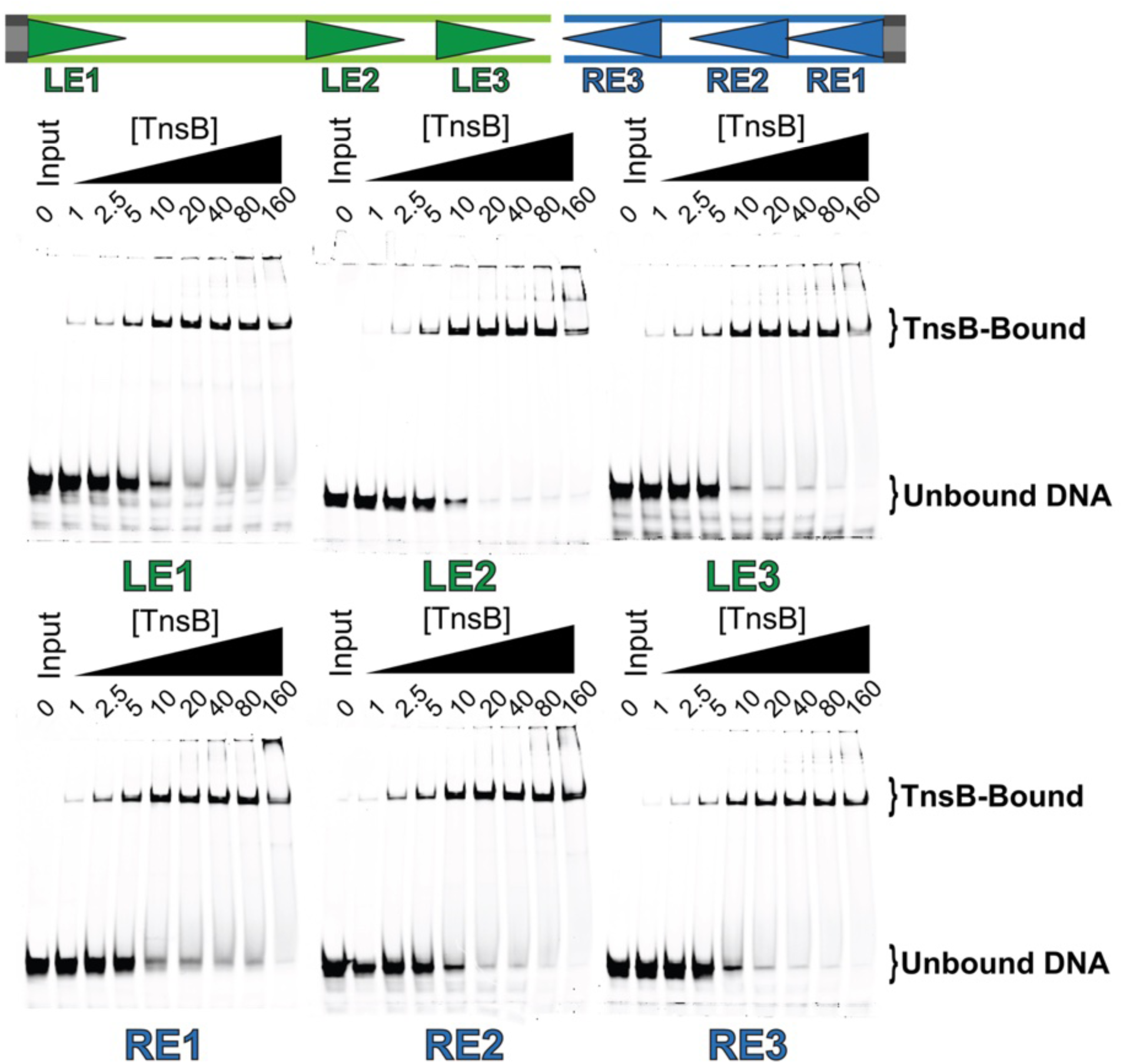
Transposase binding affinities measured across all transposase binding sites by electrophoretic mobility shift assays (EMSA). EMSA quantification of tranposase binding site affinities with TnsB. Oligos representing each individual binding site within the transposon ends are annealed and reconstituted with range of TnsB concentrations. DNA is supplied in each lane at 5 nM. Bands under the unbound DNA region represent misannealed/free DNA.

---

**Supplemental movie 1:** Architecture of the VchCAST transposase complex with synapsed ends. Three-dimensional rotational views of the cryo-EM map (threshold = 8.97) are shown, both alone and alongside the atomic model. The map is colored as in Figure 1 to distinguish different components of the complex. Results from CryoSPARC 3DFlex heterogeneity analysis are also presented. Heterogeneity maps (threshold = 0.0352) are low-pass filtered to illustrate dynamics at 8 Å resolution.

## Oligonucleotides

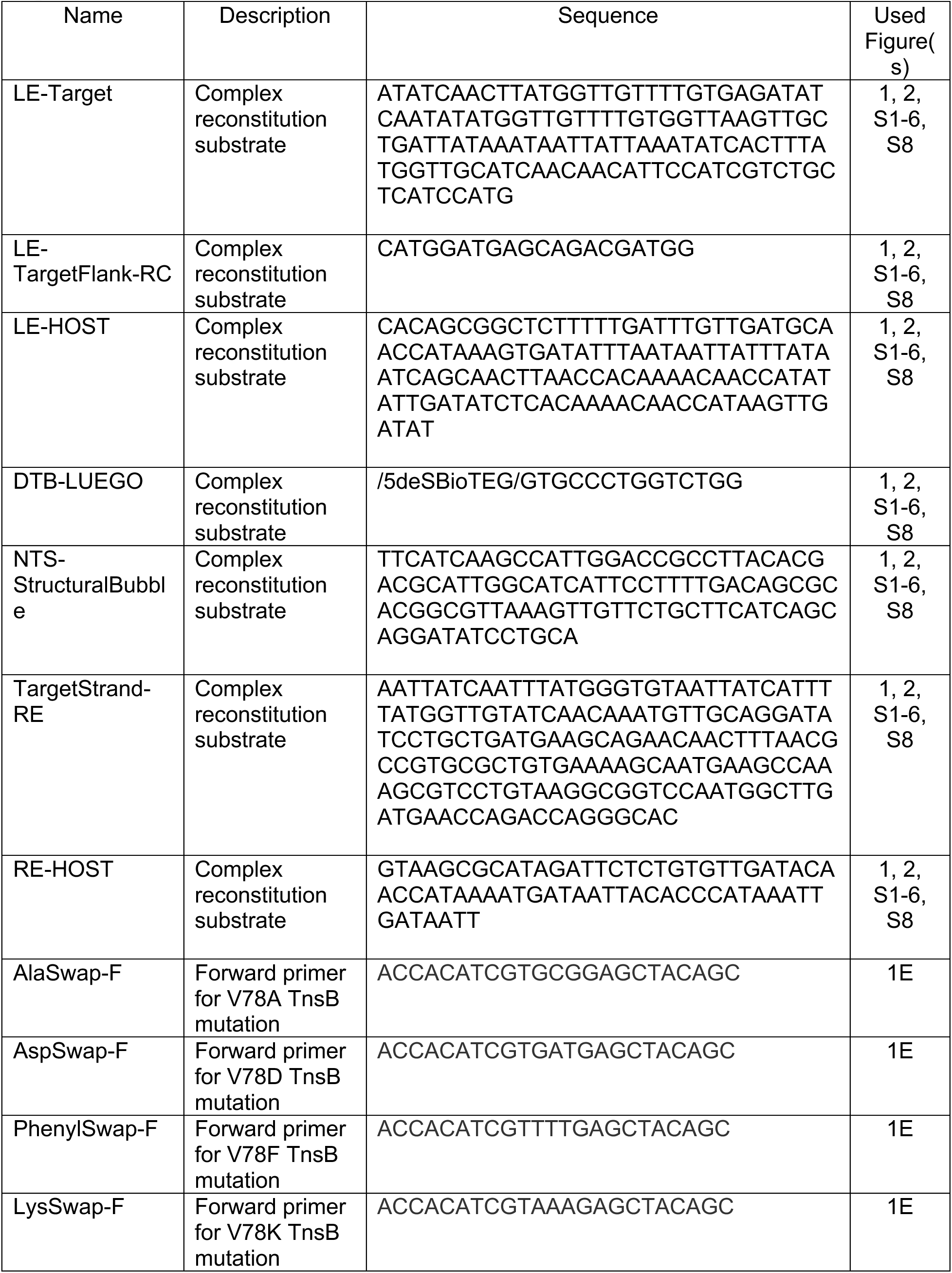
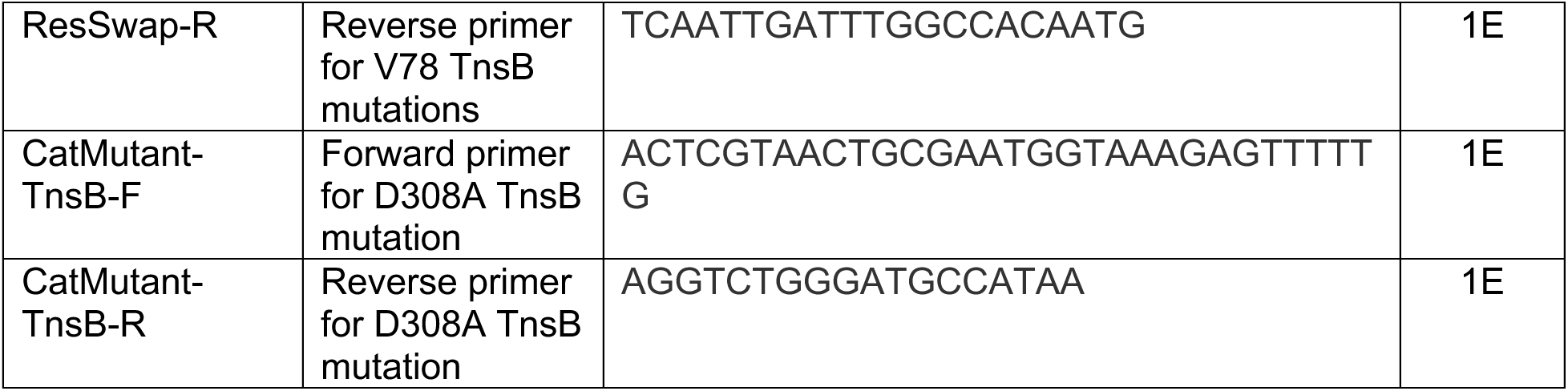

## Notes

### Competing Interest Statement

The authors have declared no competing interest.

